# Enhanced detection for antibiotic resistance genes in wastewater samples using a CRISPR-enriched metagenomic method

**DOI:** 10.1101/2024.07.30.605462

**Authors:** Yuqing Mao, Joanna L. Shisler, Thanh H. Nguyen

## Abstract

The spread of antibiotic resistance genes (ARGs) in the environment is a global public health concern. To date, over 5,000 genes have been identified to express resistance to antibiotics. ARGs are usually low in abundance for wastewater samples, making them difficult to detect. Metagenomic sequencing and quantitative polymerase chain reaction (qPCR), two conventional ARG detection methods, have low sensitivity and low throughput limitations, respectively. We developed a CRISPR-Cas9-modified next-generation sequencing (NGS) method to enrich the targeted ARGs during library preparation. The false negative and false positive of this method were determined based on a mixture of bacterial isolates with known whole-genome sequences. Low values of both false negative (2/1,208) and false positive (1/1,208) proved the method’s reliability. We compared the results obtained by this CRISPR-NGS and the conventional NGS method for six untreated wastewater samples. As compared to the ARGs detected in the same samples using the regular NGS method, the CRISPR-NGS method found up to 1,189 more ARGs and up to 61 more ARG families in low abundances, including the clinically important KPC beta-lactamase genes in the six wastewater samples collected from different sources. Compared to the regular NGS method, the CRISPR-NGS method lowered the detection limit of ARGs from the magnitude of 10^−4^ to 10^−5^ as quantified by qPCR relative abundance. The CRISPR-NGS method is promising for ARG detection in wastewater. A similar workflow can also be applied to detect other targets that are in low abundance but of high diversity.

## 1 Introduction

Antibiotic resistance (AR) is an increasing threat to global public health. It is estimated that in the next decade, if without any intervention, AR will cause an annual loss of 3.4 trillion US dollars in gross domestic products and cause 24 million more people to live in extreme poverty ^1^. In addition to surveillance in clinical settings, AR surveillance based on wastewater has been conducted ^2–9^. Culture-independent methods are commonly used for wastewater surveillance of AR ^10^, because genes that encode AR are included in bacterial genomes. By extracting and analyzing the DNA extracted from wastewater samples, the AR profiles of the entire sample can be obtained. There are two challenges to detect antibiotic resistance genes (ARGs) in wastewater DNA samples. First, the concentration of many ARGs can be very low because ARGs account for less than 1% of the DNA in bacterial genomes ^11^. Second, there is a wide range of ARG targets. To date, more than 5,000 reference sequences of ARGs are recorded in The Comprehensive Antibiotic Resistance Database (CARD) ^12^. Metagenomics and quantitative polymerase chain reaction (qPCR) are two culture-independent methods that are widely used to detect ARGs ^10^. These two methods have their pros and cons in ARG detection. qPCR has a high sensitivity, so it can capture the presence of ARGs that are in extremely low abundance in a sample ^13^. A previous study found that the detection limit of qPCR genes in manure and compost samples was 2-8 copies/mg, while that for metagenomics was around 3 × 10^4^ copies/mg ^14^. Another study also found that qPCR could stably detect ARGs in water samples, while metagenomics could not constantly detect those in low abundances ^15^. Besides, qPCR can absolutely quantify the number of gene copies in a sample ^16^. However, a major drawback of qPCR is that it only returns fluorescent signals when short DNA sequence of 70-300 bp are detected ^17^. These results can neither indicate the mutations in ARGs nor tell the existence of the remaining sequences outside of this range. This disadvantage makes qPCR easy to generate false positive results. Another drawback of qPCR is the limited number of targeted genes. Multiplexed qPCR and chip-based high-throughput qPCR microarrays are applied to increase the number of targets in one qPCR run. Multiplexed qPCR can target up to 20 genes in one reaction ^18^, but the interaction among different primers and probes may become complex when the number of targets increases and make the results inaccurate ^19^. Chip-based high-throughput qPCR microarrays can run thousands of qPCR reactions in the meantime by separating different assays into droplets, allowing it to target dozens to thousands of genes ^20^. This method can avoid the disadvantage of complex interactions of different primers and probes in conventional multiplexed qPCR while largely increasing the number of targets. However, to ensure good amplification efficiencies, the design and validation of specific primers for each individual ARG target is required. When the number of targets increases, the cost of primer design and validation will also increase. High-throughput qPCR thus has a hidden cost in the large number of primer validation. In addition, not all chip-based high-throughput qPCR machines can adjust the numbers of samples and targets flexibly ^21^. Some chip-based high-throughput qPCR machines have upper limits of 96 or 192 genes to be targeted in one run ^22^. In contrast, metagenomics does not require primer design for specific targets, so that it can detect a wide range of ARGs without pre-estimation ^23^. Next-generation sequencing (NGS) is increasingly accessible nowadays with the development of cost-friendly benchtop sequencers. However, because of the low percentage of ARGs in wastewater DNA samples, most of the reads obtained in conventional metagenomics are not relevant to ARG detection. The sensitivity of metagenomics is lower than qPCR, making the detection of low-abundance ARGs difficult by metagenomics ^10^.

To increase the sensitivity for ARG detection using metagenomics, different methods have been developed. Hybridization capture is one of the commercialized multiplexed target-enrichment methods for NGS. This method adds an additional step after indexed library preparation using streptavidin magnetic beads and specifically designed biotinylated DNA probes to capture DNA fragments with matched sequences ^24^. Thousands to millions of probes can be used to target various DNA sequences ^25^. However, this method is expensive, complex, and time-consuming because of the synthesis of highly multiplexed biotinylated DNA/RNA probes ^26,27^. Recently, CRISPR (clustered regularly interspaced short palindromic repeats)-Cas9-based target enrichment NGS methods were developed for clinical diagnosis ^28,29^. Such methods use specifically designed guide RNA (gRNA) to guide Cas9 nuclease specifically cleave the target genes to generate 5’ phosphate group for adapter ligation. The libraries generated by this method can be applied to both the Illumina NGS platform and the Nanopore 3^rd^-generation sequencing platform ^28,29^. Unlike qPCR where multiplexed primers and probes may have complex and unexpected interactions ^19^, the interaction of Cas9-gRNA complexes to DNA samples is simpler, making the Cas9 cleavage reactions possible to be highly multiplexed ^30^. Unlike biotinylated DNA probes in hybridization capture (which do not contain conserved regions), the gRNA used for Cas9 cleavage contains conserved regions ^31^, making the synthesis of multiplexed assays of CRISPR-Cas9 enriched methods more flexible and economical.

NGS is increasingly accessible to laboratories because of its reduced cost ^32^. In this study, we developed a novel CRISPR-NGS method for ARG detection in wastewater samples. The detailed step-by-step protocol for the entire CRISPR-NGS workflow can be found at <u>dx.doi.org/10.17504/protocols.io.8epv5xdnjg1b/v2</u>. This CRISPR-NGS method includes six major steps (**Figure 1**). Step 1 is to prepare a highly multiplexed gRNA pool (i.e., 6,010 different sequences) targeting different sites in ARGs specifically. We chose to synthesize trans-activating CRISPR RNA (tracrRNA) and multiplexed CRISPR RNA (crRNA) from DNA templates to reduce the cost because purchasing gRNA individually will be extremely costly ^31^. Step 2 is to prepare DNA samples for sequencing. This step includes three sub-steps of wastewater DNA extraction, DNA blocking by 5’ phosphate group removal, and DNA positive control spike. In NGS library preparation, adapter ligation is a necessary step which requires DNA fragments have 5’ phosphate group. To avoid adapters ligating to DNA fragments irrelevant to ARG detection, we used alkaline phosphatase to remove 5’ phosphate groups from all DNA fragments before library preparation. After blocking, a 250-bp DNA fragment with synthesized 5’ phosphate group obtained from *Pseudogulbenkiania* sp. strain NH8B which should be absent in most DNA samples was spiked in the DNA sample as the positive control for sequencing ^33^. Step 3 is to specifically cleave the DNA samples at the sites in the ARGs using Cas9 nuclease to generate new 5’ phosphate groups for adapter ligation. The newly generated 5’ phosphorylated ARG fragments were further modified by a dA-tailing step for optimized adapter ligation efficiency. Step 4 is to ligate the adapters to the specifically cleaved ARG fragments. Step 5 is to amplify the adapter-ligated fragments by polymerase chain reaction (PCR) using the primers targeting the conserved region on the adapters to enrich the ARG fragments in the DNA sample. Step 6 is to check the fragment lengths and the DNA concentrations of the library and load the libraries for NGS. As described here, the CRISPR-NGS method modifies the DNA fragmentation step in NGS library preparation, so that the sensitivity of NGS in ARG detection can be largely increased. With the same total number of reads, hundreds more ARGs in low abundances in sewage samples can be uncovered, and a lower detection limit can be expanded from the magnitude of 10^−4^ to 10^−5^ as determined by qPCR relative abundance.

**Figure 1.**
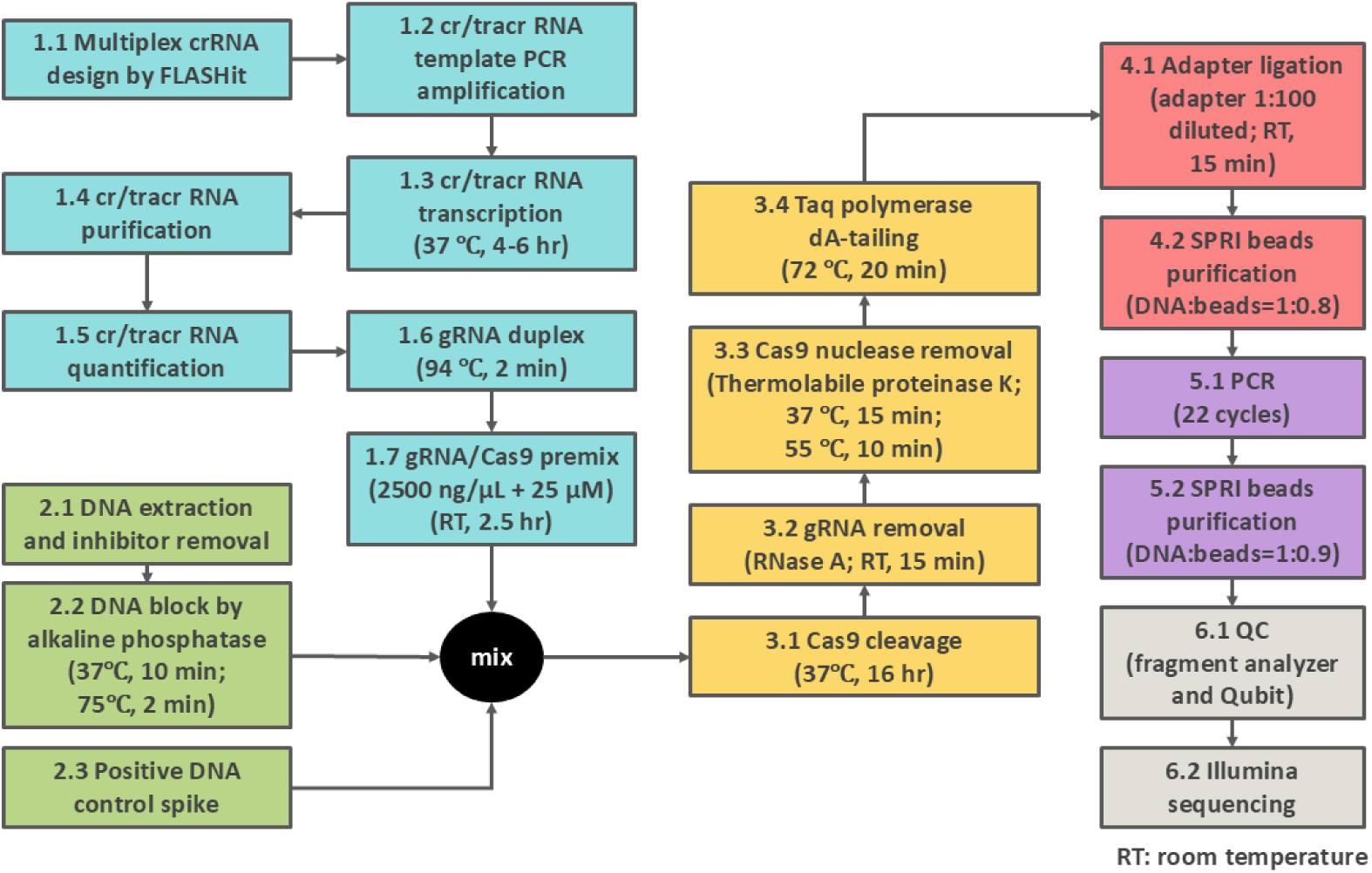
The workflow of the CRISPR-NGS method.

## 2 Methods

### 2.1 Wastewater sample collection, pre-treatment, and DNA extraction

The wastewater samples used in this study included five influents from the wastewater treatment plants in Chicago, Illinois, and one raw sewage composite samples directly collected from a manhole. More information on sample collection and processing is in **Supplementary Methods S1**.

### 2.2 Mock community DNA sample preparation

We estimated the false positive and false negative detection of ARGs using the mixture of DNA extracts at equal concentrations from 10 bacteria isolates (three *E. coli* from wastewater, three *E. coli* from clinical sources, one *Shigella* spp. from a clinical source, two *Salmonella* spp. from clinical sources, and one *Pseudomonas aeruginosa* PA14 (see **Table S1**) ^34^. This mock community was subjected to both whole-genome sequencing and CRISPR-NGS to determine the false positive and false negative rates of the CRISPR-NGS method. Details on DNA extraction, preparation, and subsequent whole-genome sequencing are in **Supplementary Methods S2**. The whole-genome sequences are available in BioProject PRJNA1147862 (to be released upon publication) ^35^.

### 2.3 Design and synthesis of crRNA and tracrRNA

Multiplexed gRNA for ARG detection was designed by FLASHit ^28^. Four thousand six hundred and forty-one ARG nucleotide sequences for gRNA design input were downloaded from the CARD protein homolog model on September 12, 2022 ^36^. The default exclusion of off-targets in the *E. coli* BL21 genome in FLASHit was removed, because the Cas9 protein used in this study was commercially purchased. In total 6,010 different 20-nt spacer sequences were generated by FLASHit to cleave target ARGs into approximately 200-bp fragments. More details of the synthesis of crRNA and tracrRNA are in **Supplementary Methods S3**.

### 2.4 CRISPR-NGS and regular NGS library preparation and sequencing

We used Cas9 nuclease to specifically cleave targeted sites for DNA fragmentation in CRISPR-NGS library preparation. For regular NGS library preparation, we used non-specific enzymes for DNA fragmentation. A synthesized 250-bp DNA fragment obtained from *Pseudogulbenkiania* sp. strain NH8B with 5’ phosphate group was spiked to all libraries as the positive control ^33^. In the adapter ligation step, xGen™ UDI-UMI Adapters (Integrated DNA Technologies) was used to reduce the PCR bias. Detailed information for CRISPR-NGS library preparation, regular NGS library preparation, library quality control, and Illumina sequencing is in **Supplementary Methods S4-S5**.

### 2.5 Sequencing read mapping

After paired-end sequencing, UMI-tools, PriceSeqFilter, Bowtie2, and SAMtools were used for read deduplication, quality control, and ARG mapping ^37–41^. The same read mapping methods were used for both CRISPR-NGS and regular NGS libraries. All sequencing raw data, including the separated UMI index files, can be found in BioProject PRJNA1148079 (to be released upon publication) ^42^. The reference file for read mapping includes the ARG sequences downloaded from CARD (see 2.3) and the sequence of the 250-bp NH8B DNA positive control. Any ARGs with coverages higher than 75% were considered present in the sample, according to previous studies ^43–45^. The identity threshold was approximately 90%, according to the end-to-end alignment algorithm of Bowtie2 ^39^. Detailed information for read mapping is in **Supplementary Methods S6**.

## 3 Results and Discussions

### 3.1 Method validation

We validated that alkaline phosphatase blocking inhibited the T4 DNA ligase from connecting different DNA fragments together. The detailed method for the validation of 5’ phosphate group removal can be found in **Supplementary Methods S7**. After phosphatase blocking, the 5’ phosphate group of DNA fragments is removed, and T4 ligase could not anneal DNA fragments to form new fragments of other sizes (**Figure S1**). We used a synthetic 731-bp DNA template with 5’ phosphorylation to test the blocking efficiency of alkaline phosphatase. After phosphatase blocking, the DNA was incubated with T4 ligase. As shown in **Figure S1**, while the DNA sample without blocking showed multiple bands, the blocked DNA remained its original 731-bp size, suggesting the 5’ phosphate groups of blocked DNA fragments were removed, and T4 DNA ligase could not ligate these DNA fragments.

We validated the Cas9 cleavage using an in-silico analysis that compares the breakpoints in raw sequencing reads with the intended Cas9 cleavage sites. In read mapping, paired 150-bp raw sequencing reads were aligned to the reference ARG sequence obtained from CARD. The paired reads, which started from different ends of the same DNA fragment, should have an overlapping region if the DNA fragment is smaller than 300 bp. If the reads are from different fragments, there would be no overlap. Because most DNA fragments were designed to be around 200 bp using a protocol developed previously ^28^ and the maximum read length was 150 bp in NGS, the entire DNA fragment would not be covered by one single read but could be covered by two paired reads from different directions. Therefore, in the read mapping plot of a cleaved 200-bp DNA fragment, we expected to see the middle part (usually around 100 bp) overlapped by the paired 150-bp reads from different directions and the remaining left and right part covered by the remaining lengths of the paired reads. Because the DNA fragments were generated by Cas9 cleavage at a few designed sites, these DNA fragments should not overlap with each other in the read mapping of the ARG. The locations where two DNA fragments meet each other should be the same as the designed Cas9 cleavage sites. The example of overlapping regions on different DNA fragments in the same ARG was marked and shown in **Figure S2,** which shows the read mapping plot for CRISPR-NGS and regular NGS for the same gene OXA-36 (ARO: 3001430) ^12^. The four cleavage sites on OXA-36 broke the read mapping histogram of CRISPR-NGS into three overlapping regions while the regular NGS read mapping histogram was smooth. We checked the read mapping of 79 ARGs from different ARG families, 78 of the raw reads were broken at the designed cleavage sites except for *rpoB* which contained a few off-target cleavage sites. This finding suggested most of the ARGs detected by CRISPR-NGS truly existed in the samples.

### 3.2 CRISPR-NGS method showed high accuracy in ARG detection in the mock community sample

We conducted whole-genome sequencing of 10 bacteria isolates (listed in **Table S1**) and these genomes served as a basis to determine false positives and false negatives in ARG detection by the CRISPR-NGS method. Because some ARGs share highly similar sequences, we divided the 4,641 targeted ARGs into 1,208 clusters based on their similarities in terms of ANI to avoid overcounting these ARGs (**Figure S3; Spreadsheet S1**). The details of ARG clustering method are in **Supplementary Methods S8**. We first used annotation for the 10 whole-genome sequences to find 100 ARGs in 99 clusters existing in the mock community. The CRISPR-NGS method detected 1,032 ARGs in 124 clusters in the same sample (**Spreadsheet S2**). Ninety-seven clusters were found by both CRISPR-NGS and annotation of the whole genomes. Because the algorithm of whole-genome annotation may miss some ARGs ^46^, we used BLAST to search the ARG sequences listed in CARD within the 10 whole bacterial genomes in the mock community ^47^. We found 25 additional clusters of ARGs that were detected by the CRISPR-NGS method in the mock community. Finally, we used Sanger sequencing to confirm the presence of two clusters of ARGs that were not found by either annotation or BLAST in the 10 whole-genome sequences, one containing quinolone resistance protein (*qnr*) genes and the other containing one single ARG *aph(3’)-Ia.* The primer sequences of these two genes are listed in **Table S2**. The amplicon of the *qnrB5* gene was aligned to the CARD reference sequence in 99.65% identity, suggesting this gene was missed by long-read PacBio sequencing but could be detected by short-read NGS. In total, as summarized in **Table S3**, we confirmed the presence of 123 ARGs cluster detected by the CRISPR-NGS method (97 from annotation of **Spreadsheet S3**, 25 from the BLAST method and one from Sanger sequencing). These clusters are the true positives. There were two false negatives, which were found by whole-genome sequencing but not by CRISPR-NGS. The maximum identities of these two ARG clusters in all mock community genomes were only 86-87% when comparing with the reference sequences, while the maximum identities of any other true positive ARGs were equal to or greater than 93%. The NGS libraries used Bowtie2 for ARG identification which has an identity threshold of approximately 90% under our current settings ^39^. In contrast, ARGs were identified from the whole-genome sequences by the Genome Annotation tool on Bacterial and Viral Bioinformatics Resource Center (BV-BRC) with an identity threshold of 80% ^36,48,49^. The difference in identity thresholds made the two ARG clusters of 86-87% identities missed by NGS.

We also compared the ARG detection results for the mock community sample to the regular NGS method using the similar methods as described above. The regular NGS library detected two more ARG clusters, *dfrA14* and *aadA13*, than the CRISPR-NGS library (**Spreadsheet S4**). However, the identities of these two ARGs were 89% in BLAST against the reference sequence. The coverage of *aadA13* (ARO: 3002613, 799 bp) in the CRISPR-NGS library was 65.91%, slightly below the 75% threshold. It is worth noting that regular NGS didn’t have any reads mapped to the reference sequence after approximately 700 bp, suggesting potential accumulation of mutations there (**Figure S4**). The cleavage site of CRISPR-NGS at 772-795 bp might be affected due to the mutations and resulted in the low coverage. The failure in *dfrA14* (ARO: 3002859, 475 bp) detection might be due to the unsuccessful cleavage at 15-38 bp (**Figure S4**). To conclude, though CRISPR-NGS may miss some ARGs with accumulated mutations that make the identities of the ARGs below 90% with reference sequences, its overall accuracy in ARG detection is high.

### 3.3 High reproducibility in ARG detection was observed in the CRISPR-NGS libraries made from the same DNA sample

The reproducibility of the CRISPR-NGS method was tested by comparing the results of two sewage samples collected from Calumet on 8/16/2023 and 10/24/2023. The detailed method is in **Supplementary Methods S9**. The DNA extracted from these samples were made into duplicated libraries. Nine hundred and sixty ARGs were detected by both libraries for Calumet (8/16/2023), with 1,020 and 1,045 ARGs detected in each library (94.1% and 91.9% were detected in both). As for Calumet (10/24/2023), 1,597 and 1,281 ARGs were detected in each duplicate, while 1,263 ARGs were shared in common (79.1% and 98.6% were detected in both).

The read depths for the ARGs detected in only one duplicated library ranged from 1.03 to 291.68, with a median of 7.70, for Calumet (8/16/2023) and ranged from 1.08 to 1041.34, with a median of 20.72, for Calumet (10/24/2023). These ARGs were missed because their coverages were below the 75% threshold, instead of no read alignment. For example, the 987-bp *AAC6_30_AAC6_Ib* gene (ARO: 3002599) detected only in one of the two libraries of Calumet (10/24/2023) with read depth of 1041.34 and coverage of 82.07%, while the other library only got a coverage of 60.59%. As a comparison, we also made duplicated regular NGS libraries for Calumet (10/24/2023). One hundred and fifty-six ARGs were detected by both libraries, while 399 and 190 ARGs were detected by each individual library (39.1% and 82.1%). The read depths of the ARGs detected only by one library ranged from 0.91 to 6.07, with a median of 1.36. These findings suggest that though having 100% same ARG detection by metagenomics is difficult, it is still possible to repeatedly detect most of the ARGs in the same DNA sample by CRISPR-NGS.

Then we compared the read depths of the ARGs detected in both duplicated libraries. The read depths of shared ARGs were fitted linearly in log-10 scale with slopes of 0.92 and 1.05, intercepts of 0.017 and −0.377 (**Figure 2**). The R^2^ values were 0.90 and 0.98, respectively, suggesting low fitting variations. The slopes of both fittings were close to 1, and the intercepts were close to 0, suggesting the read depths of the ARGs detected in the duplicated libraries were reproducible.

**Figure 2.**
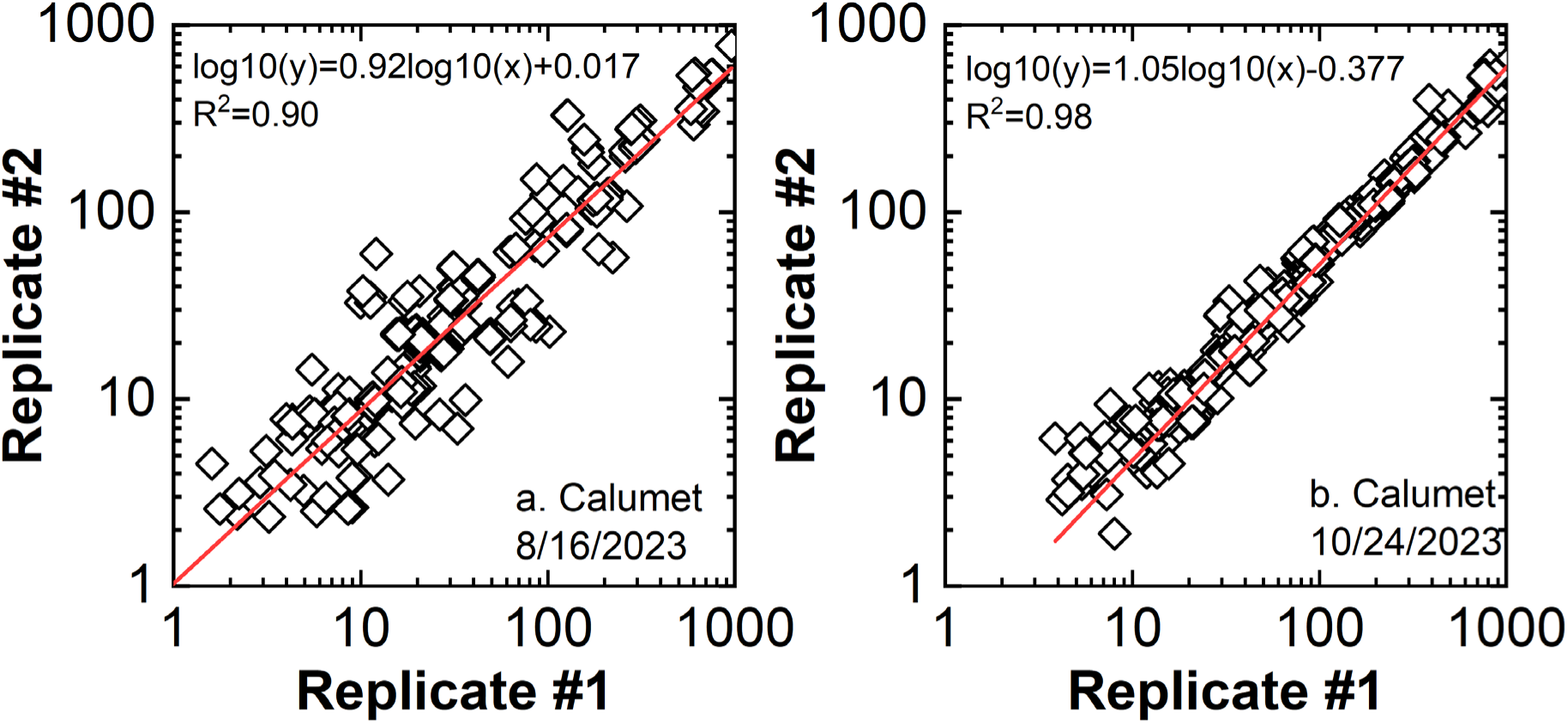
The ARG read depths in replicated libraries made from the same DNA samples (a: Calumet, 8/16/2023; b: Calumet, 10/24/2023) fitted linearly in log-10 scale with R^2^ values of 0.90 and 0.98, respectively.

### 3.4 More ARGs were detected by the CRISPR-NGS method than regular NGS method

The improved sensitivity of the CRISPR-NGS method in ARG detection was evaluated by comparing the number of ARGs and ARG families detected in the same sewage sample by the CRISPR-NGS method and regular NGS method, respectively. **Figure 3** shows the number of ARGs and ARG families that were mapped with over 75% coverage by only the CRISPR-NGS method, only the regular NGS method, and both methods. The sewage samples had 288-433 ARGs detected by regular NGS libraries, and these numbers increased to 542-1,615 for CRISPR-NGS libraries. Only from seven to 43 ARGs were uniquely detected by regular NGS libraries, while 278-1,189 ARGs were uniquely detected by CRISPR-NGS libraries. The ARGs detected in wastewater samples by different CRISPR-NGS and regular NGS libraries are listed in **Spreadsheet S5 and S6**. These additional ARGs detected by CRISPR-NGS libraries were mostly from ARG families that were not covered by regular NGS libraries. Seventeen to 61 more ARG families were only detected by CRISPR-NGS libraries for the sewage samples, while zero to three were only detected by regular NGS libraries. CRISPR-NGS libraries missed these ARG families mainly because the coverages of these ARGs were slightly lower than the 75% threshold. For example, CfxA beta-lactamase genes were missed by three CRISPR-NGS libraries (Calumet 10/24/2023, Calumet 12/19/2023, and Lemont 12/19/2023). The coverages of CfxA2 (ARO: 3003002) in these libraries were between 72.05% and 72.57%, less than 3% below the coverage threshold. The ARG families detected only by CRISPR-NGS libraries include KPC and CTX-M beta-lactamase genes which are clinically important. Another example is the mobilized colistin resistance (MCR) phosphoethanolamine transferase genes that encode resistance to colistin which is one of the last-resort antibiotics ^50,51^. The CRISPR-NGS method detected MCR genes in a high diversity, including MCR-3, MCR-4, MCR-5, MCR-9, and MCR-10, while regular NGS only detected MCR-3 or MCR-5 in some libraries.

**Figure 3.**
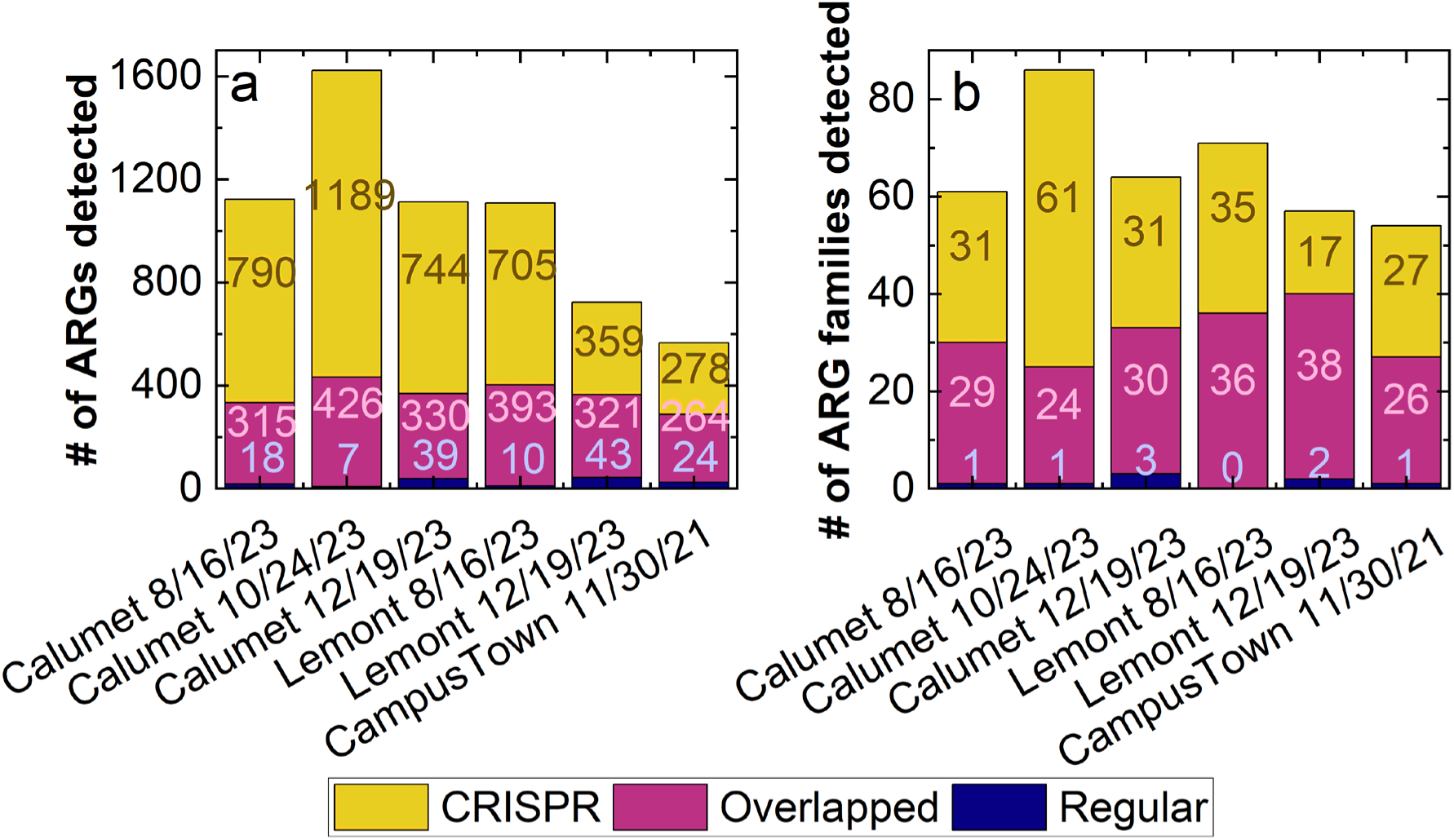
More ARGs (a) and ARG families (b) were detected by CRISPR-NGS method than regular NGS method in different sewage samples. The yellow bars show the number of ARGs and ARG families only detected by CRISPR-NGS method, the purple bars show the number of ARGs and ARG families detected by both CRISPR-NGS and regular NGS methods, and the dark blue bars show the number of ARGs and ARG families only detected by the regular NGS method.

### 3.5 CRISPR-NGS method could detect ARGs in higher read depths than regular NGS method

The ARGs detected by CRISPR-NGS libraries also had higher read depths than those detected by regular NGS libraries. The fold changes in read depths of the 145 ARGs detected in all sewage samples by both CRISPR and regular NGS libraries are plotted in **Figure 4**. All ARGs had significantly higher read depths in CRISPR-NGS libraries than regular NGS libraries. The fold changes in read depths varied from 2.73 to 126.45, suggesting the CRISPR-NGS method can detect ARGs in higher confidence than regular NGS method.

**Figure 4.**
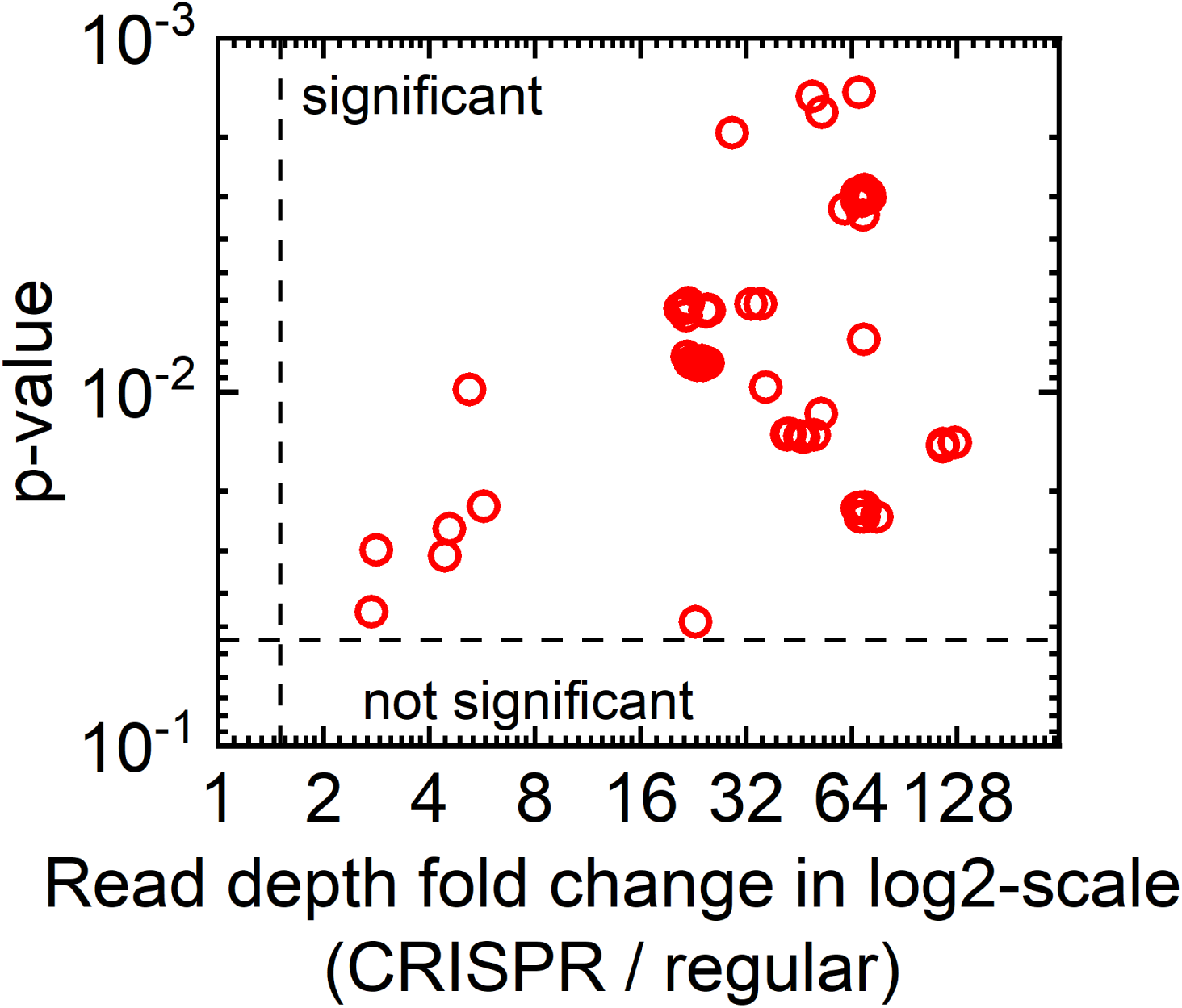
The read depths of all 145 ARGs detected in all sewage samples were significantly higher in CRISPR-NGS libraries than regular NGS libraries. The y axis shows the p-values in a reversed log-10 manner, and the x axis shows the fold changes in read depths (CRISPR-NGS versus regular NGS). The horizontal dash line means the threshold of a significant p-value equals to 0.05, and the vertical dash line means the thresholds in fold change of 1.5.

To compare the ARG detection efficiencies of CRISPR-NGS and regular NGS with deeper sequencing read depths, we subsampled the CRISPR-NGS reads and compared the ARG mapping results with the corresponding regular NGS libraries without subsampling (**Figure 5**). The subsampling protocol can be found in **Supplementary Methods S10**. Before subsampling, the total number of reads for CRISPR-NGS libraries for sewage samples ranged from 43.3 million to 58.2 million, and the total number of reads for regular NGS libraries ranged from 44.6 million to 60.7 million (**Table S4**). When subsampling 2% of the reads, three CRISPR-NGS libraries (Calumet 12/19/2023, Calumet 10/24/2023 A and B) detected more ARGs than the corresponding regular NGS libraries. When the subsampling ratio reached 10%, all CRISPR-NGS libraries except for Lemont (12/19/2023) detected more ARGs than corresponding regular NGS libraries. The CRISPR-NGS library for Lemont (12/19/2023) detected 89.6% of the number of ARGs detected by its regular NGS library when doing 10% subsampling, and the percentage increased to 99.2% at subsampling ratio of 20%, and 171.2% at subsampling ratio of 50%. At subsampling ratio of 50%, all curves reached a plateau phase (**Figure 5**). These findings suggest that CRISPR-NGS can detect similar numbers of ARGs as regular NGS with approximately 2-20% of the total number of reads. With approximately 25 million total number of reads (i.e., approximately 50% of the read depths used in this study), the number of ARGs detected by CRISPR-NGS approaches maximum. This improvement suggests that CRISPR-NGS can make it possible to detect ARGs deeply by metagenomics on many benchtop sequencers that can generate more than 25 million reads in one run, instead of ultra-deep sequencing for regular NGS libraries that usually require expensive and professional production-scale sequencers.

**Figure 5.**
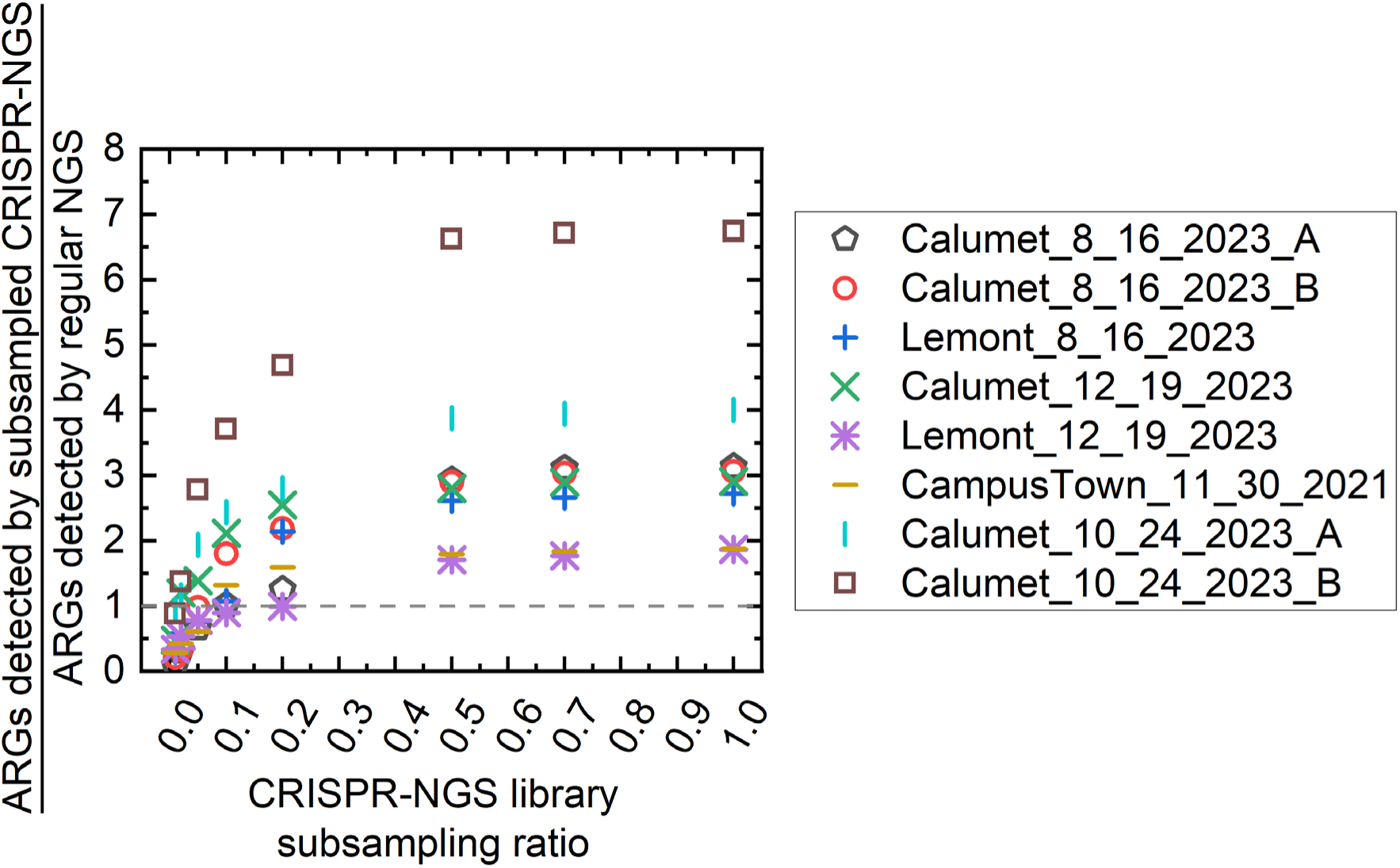
When the total read depth reaches approximately 2-20% of current read depth, CRISPR-NGS libraries can detect similar numbers of ARGs as regular NGS libraries with their original read depths. The CRISPR-NGS libraries were subsampled by ratios of 1%, 2%, 5%, 10%, 20%, 50%, and 70%. The horizontal dash line at y=1 means equal number of ARGs detected by the subsampled CRISPR-NGS library and the regular NGS library with original read depth.

### 3.6 ARG presence could be validated by Sanger sequencing

We used Sanger sequencing to validate the presence of selected ARGs in sewage samples (**Table 1**). The ARG targets were amplified by PCR, and the PCR amplicons were Sanger sequenced and compared to the reference sequences in CARD. The detailed method for Sanger sequencing validation can be found in **Supplementary Methods S11**. We chose Sanger sequencing for validation, because Sanger sequencing for PCR amplicons is accurate, cost-effective, and fast when there are only a few targets. The identities between the sequenced PCR amplicons and the reference sequence of the ARGs in CARD were between 89% and 100%, suggesting the amplicon sequences were highly similar to the reference sequences. Because the PCR primers designed for different ARGs had different binding regions, the amplicons could not cover the full lengths of the ARGs. After adjusting with the amplicon lengths, the coverage of the ARGs were between 58% and 99%. The 58% coverage of *aac(3)-IId* and 73% coverage of *aph(3’)-Ia* were due to the low quality of Sanger sequencing of the campus town manhole sample. After removing these outliers, the coverages were between 77% and 99%. The high coverages and identities of Sanger sequencing also matched the read mapping plots. For example, the two colistin resistance genes, MCR-5.1 and MCR-4.1, were nearly fully mapped by CRISPR-NGS reads (**Figure S5**), though these two genes were barely detected by regular NGS.

**Table 1.** The Sanger sequencing identities and adjusted coverages for the ARGs detected in different CRISPR-NGS libraries.

| Gene | Identity (%) | Coverage adjusted by amplicon lengths (%) |
| --- | --- | --- |
| *aph(3'')-Ib* | 99-100 | 95-98 |
| *sul1* | 97-99 | 95-96 |
| *CRP* | 89-94 | 95 |
| *sul2* | 95-99 | 92-96 |
| *aph(3')-Ia* | 95-99 | 73-97 |
| *dfrA1* | 96-97 | 93-95 |
| *GES-26* | 95-100 | 96-97 |
| *MCR-5.1* | 98 | 77-99 |
| *MCR-4.1* | 98 | 98 |
| *TEM-1* | 99 | 96 |
| *aac(3)-IId* | 96-99 | 58-93 |
| *tet(G)* | 94 | 88 |
| *qnrB5* | 100 | 97 |

**Note:** The coverages were adjusted by dividing the aligned length by the amplicon length, rather than the coverage provided in BLAST, which divides the aligned length by the query length (i.e., the reference ARG length in CARD).

### 3.7 CRISPR-NGS libraries had lower detection limit than regular NGS libraries, and the read depths in CRISPR-NGS libraries were proportional to the relative abundances determined by qPCR

We evaluated the detection limit of the CRISPR-NGS method by plotting the read depths against the relative abundance detected by qPCR for nine ARGs with various read depths detected by CRISPR-NGS or regular NGS methods in sewage samples (**Figure 6**). The detailed method for qPCR can be found in **Supplementary Methods S12**. The ARGs and corresponding qPCR primers are listed in **Table S5**. The Cq values of the 16S rRNA gene in the sewage samples are listed in **Table S6**. The read depths in both CRISPR-NGS and regular NGS libraries increased when the qPCR relative abundances increased, suggesting the read depths of CRISPR-NGS and regular NGS can be used for ARG abundance estimation. As shown in **Figure 6**, the lower detection limit of the CRISPR-NGS method for these nine ARGs was down to 2.23×10^-5^ copies ARG per copy of 16S rRNA gene, while the lower detection limit of regular NGS method was at 1.79×10^-4^ copies ARG per copy of 16S rRNA gene. Thus, the CRISPR-NGS method has a higher sensitivity than regular NGS method in ARG detection. A previous study reported varied relative abundances of ARGs in wastewater samples ranging from 1.4×10^-5^ to 0.09 ^52^, suggesting the improved lower detection limit of CRISPR-NGS can promote the detection of ARGs in low abundances in wastewater samples.

**Figure 6.**
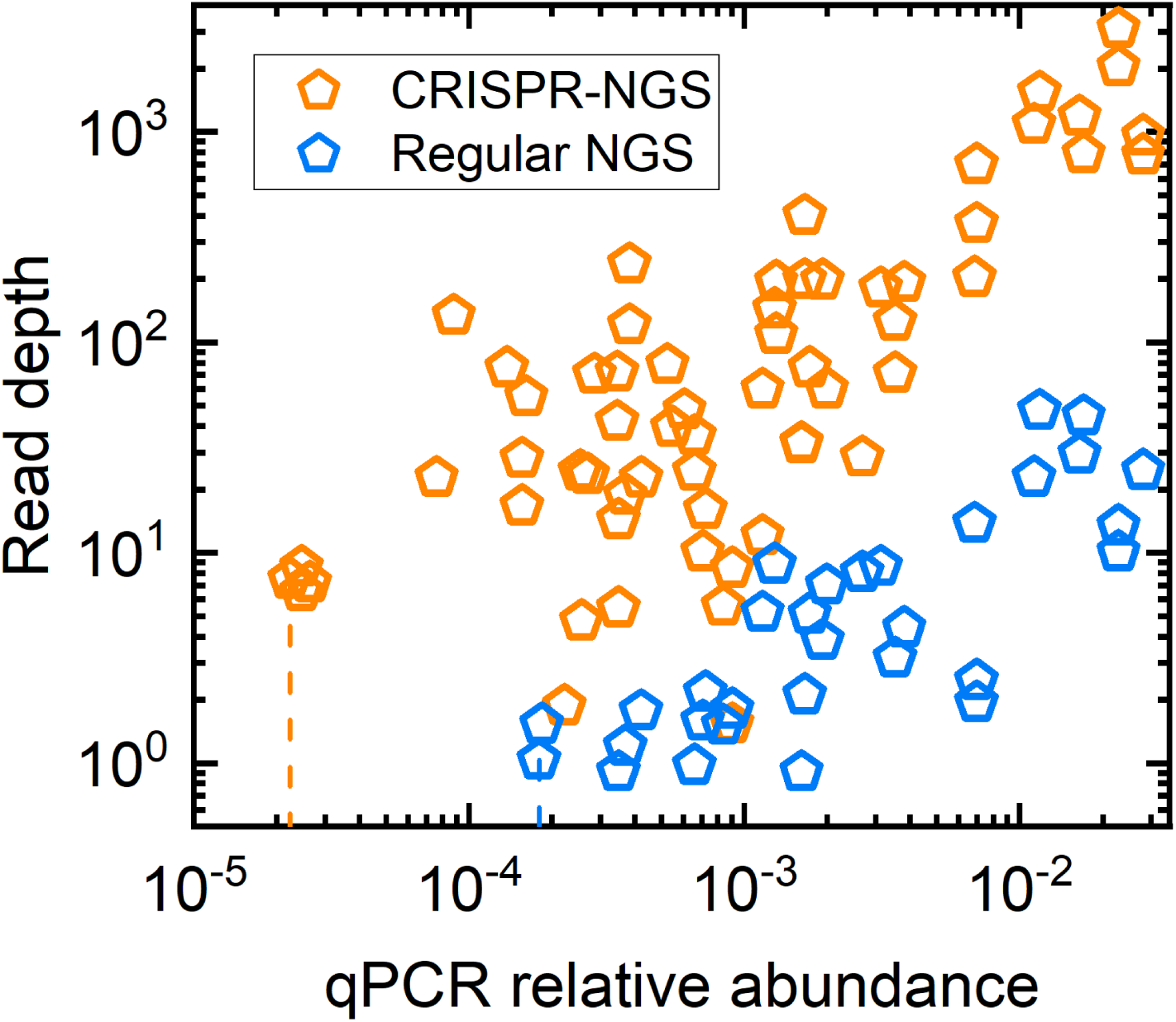
The read depths versus the qPCR relative abundances of the nine ARGs detected in different wastewater samples. The orange dots are the read depths of the ARGs detected in the CRISPR-NGS libraries. The blue dots are the read depths of the ARGs detected in the regular NGS libraries. The vertical dash lines in corresponding colors indicate the lower detection limits of CRISPR-NGS and regular NGS in terms of qPCR relative abundance. CRISPR-NGS had a lower detection limit of 2.23×10^-5^, as labeled by the orange vertical line, while the lower detection limit of regular NGS was 1.79×10^-4^, as labeled by the blue vertical line.

### 3.8 CRISPR-NGS is better at detecting low abundance ARG than regular NGS

The most abundant clusters of ARGs (i.e., ARG clusters with read depths greater than 200 in any of the CRISPR-NGS libraries) in the sewage samples were selected and plotted into heatmaps for comparison in **Figure 7**. These 41 ARG clusters were distributed in eight major drug classes, including aminoglycoside (8), beta-lactam (17), antiseptics (2), fluoroquinolone (2), macrolide (3), multi-drug (3), phenicol (2), and tetracycline (4). The ARG clusters in **Figure 7** are named based on the ARG names and families in the cluster. The multi-drug class was defined to include the ARGs that are resistant to at least two drug classes in the 61 sub-terms under the CARD “antibiotic molecule” ontology term ^12^. Comparing between CRISPR-NGS and regular NGS libraries, the entire group of CMY-2 and KPC beta-lactamase genes and *qnrB8*-like quinolone resistance genes were missed by regular NGS. Comparing the wastewater treatment plant sewage samples, the campus town manhole sample had lower abundances of GES, MOX-9, and OXA-427-like beta-lactamase genes than the wastewater treatment plant (WWTP) influent samples. The abundance of the *tet(C)* gene in the campus town manhole sample was greater than any of the Chicago WWTP influent samples, but *tet(E)* in the campus town manhole sample was lower than the WWTP influent samples. The above findings suggested different ARG distributions in different geographical areas or different sampling scales. As for the ARGs shared in common, *sul1*, *qacE/qacEdelta1*, *qacL*, and OXA-2-like beta-lactamase genes were constantly more abundant than the other ARGs in all sewage samples. Since *sul1*, *qacE*, and *qacEdelta1* are known to be located on class 1 integrons which play an important role in the spread of ARGs, our findings suggest that class 1 integron may be actively transferring ARGs in these wastewater samples ^53^.

**Figure 7.**
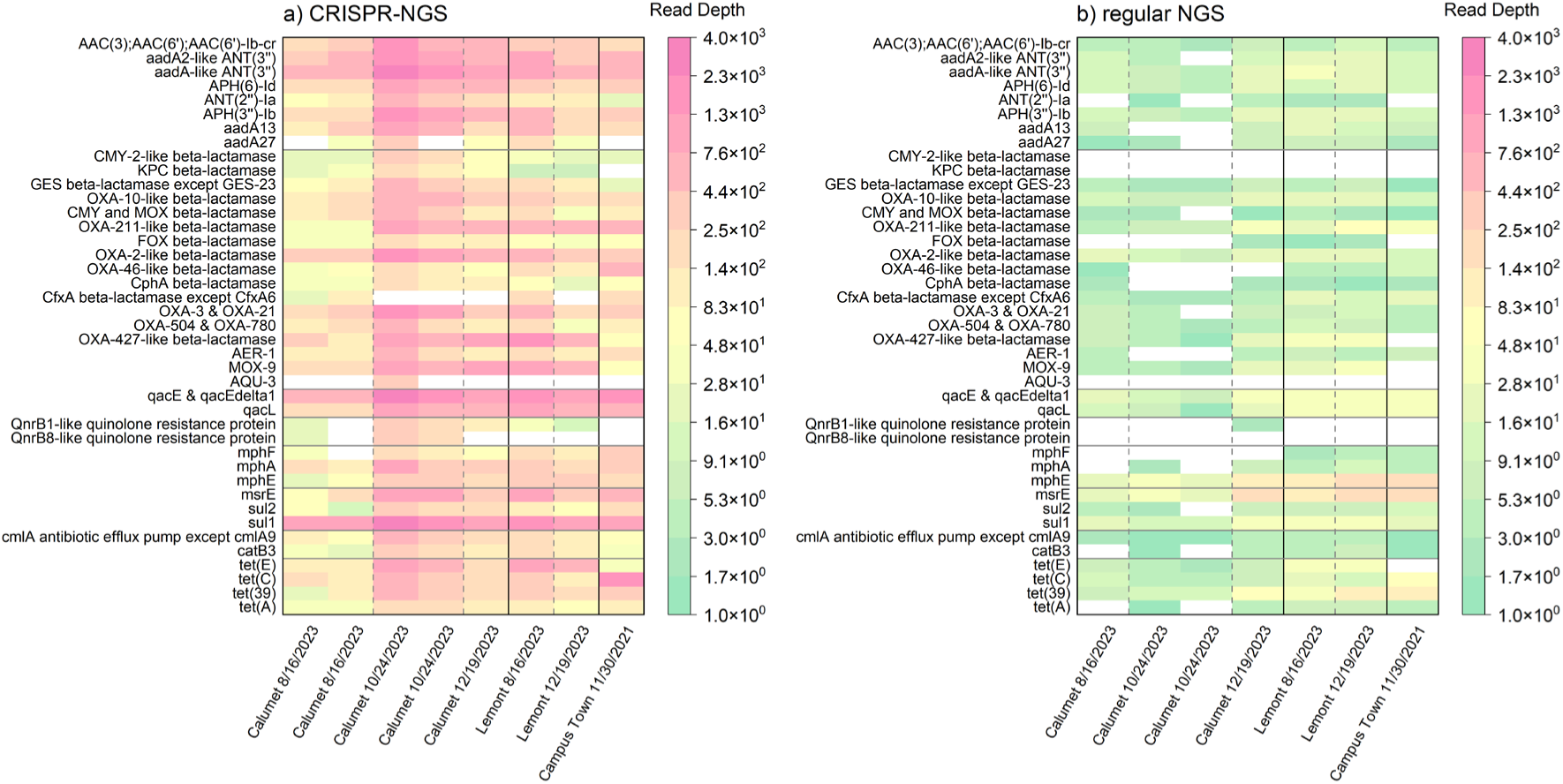
The heatmaps for the most abundant ARGs (i.e., the ARGs with read depths greater than 200 in at least one library detected by CRISPR-NGS) in different sewage samples. a: The read depths of the most abundant ARGs detected by CRISPR-NGS libraries; b: The read depths of the most abundant ARGs detected by regular NGS libraries. The read depths increase from green to pink. The white color means the ARG was not detected in the sample by the corresponding method. From top to bottom, the heatmaps were divided into eight parts by horizontal black lines according to the drug classes of the corresponding ARGs: aminoglycoside, beta-lactam, antiseptic, fluoroquinolone, macrolide, multi-drug, phenicol, and tetracycline. The solid vertical lines divide samples by different locations of collection, and the dash vertical lines divide samples from the same location by different collection times.

When extending the scale to all ARGs with high or low abundances, the profiles detected by CRISPR-NGS and regular NGS were more different (**Figure S6**). The entire class of aminocoumarin antibiotics was not detected by regular NGS libraries but by CRISPR-NGS libraries. Fewer classes of beta-lactamase genes were detected by regular NGS libraries than CRISPR-NGS libraries. For example, KPC and CTX-M were not detected by any regular NGS libraries but by CRISPR-NGS libraries. Similarly, regular NGS libraries missed several clusters of multi-drug, peptide, diaminopyrimidine, and phenicol resistance genes which were captured by CRISPR-NGS libraries. But for aminoglycoside, macrolide, and tetracycline resistance genes, regular NGS libraries detected similar numbers as CRISPR-NGS libraries. Such findings suggest that when the range of ARGs for surveillance expands, CRISPR-NGS works better in capturing the ARGs in the classes that are less abundant than the three drug classes mentioned above.

## 4 Discussion

Metagenomics has been applied widely in ARG surveillance in different water and wastewater samples ^54–58^. The advantage of metagenomics is that it can detect a wide range of ARGs without knowing the targets. However, it is challenging to sequence ARGs present in low abundances by metagenomics. We overcome this disadvantage by using CRISPR-Cas9 with multiplexed gRNA to specifically enrich the ARGs in wastewater DNA samples during NGS library preparation.

After the CRISPR-Cas9 treatment, compared to regular NGS libraries, up to 1,189 more ARGs were detected, and CRISPR-NGS libraries only require 2-20% total number of reads to detect similar number of ARGs as regular NGS libraries. Based on the findings in this study, CRISPR-NGS works the best for samples that contain highly diverse ARGs in relatively low abundances. Various types of extended-spectrum beta-lactamase (ESBL) genes were detected in the sewage samples in this study by CRISPR-NGS. These genes include not only the common ones like TEM, CTX-M, and OXA, but also the rare ones, such as GES, PER, VEB, BES, and TLA ^59^. Among these genes, GES, PER, OXA, and VEB were abundant enough to be constantly detected by regular NGS. Other genes were too diluted to be stably detected without CRISPR treatment before NGS, including the clinically important carbapenem resistance gene KPC and cefotaxime resistance gene CTX-M. The return is low but extra effort is required to monitor these rare ESBLs using conventional methods because their abundances were too low to be detected in most cases. However, studies showed that these ESBLs were also evolving and spreading globally ^60,61^. For example, GES-2 gained resistance to carbapenem by only a single point mutation ^62^. Though rarely detected in the clinical isolates obtained until now, our findings indicated that these ESBL genes were circulating in human communities and could potentially cause public health problems in the future. For example, though considered as a less common ESBL, the read depths of GES in WWTP samples were in the top 17 of all 76 detectable beta-lactamase gene clusters. In contrast, GES was not as abundant as in WWTP samples for the campus town manhole sample. This finding suggested that the presence and abundance of different ESBL may be dynamic. Innovative high-throughput detection methods with high sensitivity such as CRISPR-NGS will be helpful in the surveillance of the emergence of such unexpected ARGs.

The CRISPR-NGS method is promising for ARG detection, especially for the surveillance of sewage samples which may have many unexpected targets. Early warning of new threats in public health is challenging, because lots of new targets may be hidden in low abundance in human communities and suddenly emerge to cause an outbreak ^63^. Surveillance methods should be high in throughput and sensitivity simultaneously. When the throughput is high, multiplexed or chip-based qPCR is labor-intensive in primer design and sample preparation. Because of such reasons, NGS is more suitable for monitoring unexpected targets than qPCR. Conventional NGS can increase its sensitivity by increasing the total number of reads during a sequencing run ^64,65^. However, this method can be costly, because increasing the total read depth by 10-fold means the total sequencing cost for the same sample is increased by 10-fold. Currently, the Illumina sequencing cost is around $2.2 per million reads ^66^. For a library with 50 million reads, the cost solely for sequencing is around $110. If the total number of reads increases by 10-fold to 500 million, the sequencing cost for the same library will increase to around $1,100. Considering more than 99.9% reads are irrelevant to ARG detection ^24^, increasing the proportion of ARG reads before loading the samples to sequencing machine will be more economical. The CRISPR-NGS method could detect the same number of ARGs read depth using 2-20% of total number of reads required by regular NGS libraries with only an increased cost of around $30 in Cas9 nuclease and gRNA preparation for each library. The relatively low cost makes the CRISPR-NGS method attractive for routine ARG surveillance in the future. The reduced read depth requirement of CRISPR-NGS make it possible to use downgraded but less expensive benchtop sequencers like MiSeq to detect ARGs by metagenomic sequencing and generate results comparable to the more advanced production-scale sequencers like NovaSeq.

There are also limitations of the CRISPR-NGS method for future improvement. First, genetic information other than ARGs, for example, the pathogen genomes and mobile genetic elements, may be missed by CRISPR-NGS method, because these sequences are not targeted during assay design. ARGs that are currently unknown may also be missed for the same reason. To detect these targets, future studies can include the sequences of genetic markers of pathogens and mobile genetic elements in gRNA design. Combining CRISPR with 3^rd^ generation long-read sequencing can also help identify the host information of ARGs, because the upstream and downstream sequences of target ARGs can be identified accordingly ^67^. Second, the read depths of ARGs may be biased because of the high number of PCR cycles used during CRISPR-NGS library preparation if using conventional adapters. UDI-UMI adapters with molecule-specific adapter sequences are recommended to reduce the PCR bias, because the over-amplified DNA fragments can be identified by their shared UMI indexes. Third, CRISPR-NGS cannot quantify ARGs in absolute copy numbers as qPCR, the relative quantification is also not as accurate as qPCR. We recommend using CRISPR-NGS periodically for wastewater surveillance to identify ARGs relevant for public health and use qPCR to quantify these ARGs in a more frequent manner. Fourth, between seven to 43 ARGs were missed by CRISPR-NGS but captured by regular NGS in sewage samples, suggesting not all ARGs were successfully targeted by CRISPR-NGS. This may be due to the low on-target cleavage efficiency of some gRNA or due to “dropouts” in template DNA oligo pool synthesis. Though Cas9 nuclease can cleave DNA at targeted sites with a few mismatches ^68^, the efficiency of Cas9 binding and cleavage is reduced when mismatches occur. In addition, the current DNA oligo pool synthesis technique cannot guarantee 100% synthesis of all oligos. Around 0.8% of the sequences in a DNA oligo pool can be missed during synthesis. For the oligo pool used in this study with 6,010 different sequences, the number of “dropout” oligos is around 48. The corresponding gRNA would not be transcribed and used in library preparation. Selecting a vendor with relatively lower “dropout” ratio for oligo pool synthesis may help increase the on-target cleavage ratio. Fifth, the CRISPR-NGS method may require a more flexible read mapping threshold (e.g., different coverage threshold for different ARGs) to avoid false negative results, making the bioinformatic pipeline after sequencing more complex. In this study, we used a fixed coverage threshold of 75% to make CRISPR-NGS results comparable to regular NGS, but future studies can develop a bioinformatic pipeline with flexible threshold specifically for CRISPR-NGS. Because of the recognition of the protospacer adjacent motif (PAM) regions, gRNA sequences cannot be designed exactly at the beginning and the end of the ARGs. This disadvantage in gRNA design can result in reduced coverage in read mapping of some ARGs, causing false negative results of these ARGs when using a fixed threshold.

## 5 Conclusions

Here, we introduced a CRISPR-Cas9 modified NGS library preparation method to increase the sensitivity in ARG detection in sewage samples. Up to 61 ARG families or up to 1,189 ARGs present in low abundance were uncovered by NGS after CRISPR-Cas9 modification. With the same total read depth for one sequencing run, CRISPR-NGS libraries lower the detection limit from magnitude of 10^-4^ to 10^-5^ as determined by qPCR relative abundance. CRISPR-NGS could detect similar number of ARGs as regular NGS with only 2-20% required total number of reads. CRISPR-NGS only requires approximately 25 million reads to detect ARGs deeply in sewage samples, making it possible to be applied to cost-friendly and more accessible benchtop sequencers.

## Acknowledgement

We acknowledge funding from Water Research Foundation (5182) and USEPA (R840487). This study has not been formally reviewed by EPA. EPA does not endorse any products or commercial services mentioned in this publication. Our study was supported by Dr. Wang, Akin, and Dwivedy (CRISPR protocol), Schmidt, McAllister, Craft, Hoppenworth (sample collection and processing), Drs Whitaker, Chandrashekhar, and Suttenfield (*Pseudomonas aeruginosa* PA14). Roy J. Carver Biotechnology Center in their sequencing services, Drs Ceniceros, Frederick, and Wellman from Carle Health for providing clinical bacteria isolates, Drs. Cox, El-Naggar, Patel, and Patterson from Metropolitan Water Reclamation District of Greater Chicago for providing WWTP sewage samples.

## Supplementary Information

### Supplementary Methods

#### S1. Wastewater sample collection, pre-treatment, and DNA extraction

The wastewater treatment plant influents were taken as 24-hour composites on weekdays. The manhole sewage sample was collected by an autosampler programmed to collect four-day composite sewage samples at four-hour intervals as described in a previous study (Oh et al., 2022). After collection, the water samples were sealed in sterile sample bags and kept in coolers filled with ice packs during transportation and stored at 4 ℃ for fewer than three days before filtering. Detailed information of the samples is listed in **Table S7**. Wastewater samples were shaken well and filtered through 47 mm mixed cellulose esters (MCE) membrane filters. After filtering, the filters were cut into quarters and stored at −80 ℃ before DNA extraction. DNA was extracted from each filter using the FastDNA SPIN Kit for Soil (MP Biomedicals), using the protocol provided by the manufacturer. The centrifugation step after bead beating was increased to 15 minutes to ensure a thorough separation of liquid and solid phases. The DNA extracts were eluted by 100 μL DES solution provided in the kit. The DNA samples were purified by OneStep PCR Inhibitor Removal Kit (Zymo Research). The DNA yield was determined by Qubit™ 1X dsDNA High Sensitivity (HS) Assay Kit (Invitrogen). Then, the DNA samples were stored at −20 ℃ before library preparation.

#### S2. Mock community DNA sample preparation and whole-genome sequencing

ZymoBIOMICS DNA Miniprep Kit (Zymo Research) was used for DNA extraction. Before DNA extraction, each bacterial strain was purified by streaking onto culture plates from their frozen stocks and incubated at 37 ℃ for 24 hours. Then, single colonies were picked from the plates and spiked into 900 μL of LB broth and incubated at 37 ℃ overnight for bacteria enrichment. The overnight cultures were centrifuged at 600x g for 5 minutes to separate bacteria cells from the liquid medium. The pellet was re-suspended by 750 μL of ZymoBIOMICS Lysis Solution and directly transferred to ZR BashingBead™ Lysis Tubes. Further DNA extraction occurred by following the manufacturer’s protocol. The DNA was eluted in 100 μL of ZymoBIOMICS™ DNase/RNase Free Water. The DNA yield was determined by Qubit™ 1X dsDNA High Sensitivity (HS) Assay Kit (Invitrogen) immediately after extraction. After mixing the DNA extracts of the 10 isolates, the mock community was stored at −20 ℃ before library preparation.

GenFind V3 (Beckman Coulter) was used to extract high molecular weight DNA from each isolate listed on **Table S1**. These DNA extracts were subjected to PacBio HiFi whole-genome sequencing. Similar to above, a portion of frozen bacterial stock was streaked for purification and incubated in liquid medium for enrichment using the same method as mentioned in section 2.2. After the overnight incubation at 37 ℃ of the liquid culture, an additional enrichment step was performed by transferring 100 μL of the overnight culture into 900 μL fresh LB broth and incubating at 37 ℃ for 2-3 hours to ensure the integrity of the genome DNA. Each DNA sample was eluted by 200 μL 10 mM Tris-HCl (pH=8). The concentrations and the lengths of the DNA extracts were determined by Qubit™ 1X dsDNA High Sensitivity (HS) Assay Kit (Invitrogen) and 1% agarose gel electrophoresis. The DNA extracts were stored at −80 ℃ before further analysis.

Before PacBio HiFi library preparation, all DNA extracts were sheared by Megarupter 3 (Diagenode) at speed 40 to obtain mean fragment lengths at 10 kbp for each sample. After shearing, the sheared DNA samples were purified by OneStep PCR Inhibitor Removal Kit (Zymo Research) followed by 1:1 beads cleaning using SMRTbell® cleanup beads (PacBio). The fragment lengths of the sheared DNA samples were determined by 1% agarose gel electrophoresis. Libraries were made using the SMRTbell prep kit 3.0 (PacBio). Qubit™ 1X dsDNA High Sensitivity (HS) Assay Kit (Invitrogen) was used to determine the concentrations of the sheared DNA samples and the libraries. The fragment lengths of the libraries were determined by AATI Fragment Analyzer by Roy J. Carver Biotechnology Center before sequencing. Sequencing was also conducted by Roy J. Carver Biotechnology Center on SMRT Revio cell (PacBio) with 30-hour movie time.

The HiFi reads of the bacteria isolates were assembled into whole genomes by Trycycler ^1^ using Canu, Flye, and Raven as the assemblers ^2–4^. Then, the fasta files containing all assembled contigs were uploaded to Bacterial and Viral Bioinformatics Resource Center (BV-BRC) and annotated by the Genome Annotation service ^5,6^. The records of the ARGs identified in these genomes from the CARD source were downloaded and labeled by their corresponding AROs using a customized Python code ^7^. Due to the difference in the IDs of the ARGs used in BV-BRC Genome Annotation and CARD downloadable records, the remaining ARGs that were not automatically matched by the customized code were BLASTed manually to compare the sequences and labeled by corresponding AROs if the identity and coverage of the BLAST result were both greater than 95% ^8^. The AROs of the ARGs that were not used for crRNA design were excluded. This list of AROs was used as a reference to determine the true/false positive/negative in ARG detection for CRISPR-NGS method.

#### S3. Design and synthesis of crRNA and tracrRNA

Multiplexed gRNA for ARG detection was designed by FLASHit ^9^. Four thousand six hundred and forty-one ARG nucleotide sequences for gRNA design input were downloaded from the CARD protein homolog model on September 12, 2022 ^7^. The default exclusion of off-targets in the *E. coli* BL21 genome in FLASHit was removed, because the Cas9 protein used in this study was commercially purchased. In total 6,010 different 20-nt spacer sequences were generated by FLASHit to cleave target ARGs into approximately 200-bp fragments.

For financial reasons, multiplexed gRNA was assembled using crRNA and tracrRNA that were transcribed separately from corresponding DNA templates. The template for tracrRNA was a double-stranded synthesized DNA fragment (5’-AGGCGAATCAGATAATCGTTATGTCCAGACTGTATTAATACGACTCACTATAGGACAGC ATAGCAAGTTAAAATAAGGCTAGTCCGTTATCAACTTGAAAAAGTGGCACCGAGTCGGTGCTTTTT-3’) ^9^, and the template for multiplex crRNA was a single-stranded DNA oligo pool of the following format (The 20 Ns correspond to the 20-nt spacer sequences generated by FLASHit): 5’-TAATACGACTCACTATAGNNNNNNNNNNNNNNNNNNNNGTTTTAGAGCTATGCTGTTTTG-3’. The 20-nt spacer sequences in crRNA used in this study and their starting and ending positions in all ARGs are listed in **Spreadsheet S7**.

A PCR step was performed to enrich both templates before transcription. Each PCR reaction included 25 μL of Phusion® High-Fidelity PCR Master Mix with HF Buffer (New England Biolabs), 1.25 μL of 10 mM forward (cr/tracr: 5’-TAATACGACTCACTATAG-3’) and reverse (cr: 5’-CAAAACAGCATAGCTCTAAAAC-3’; tracr: 5’-AAAAGCACCGACTCGGTGCCAC-3’) primers respectively, 4 μL of 10 ng/μL multiplexed crRNA template or 1 ng/μL tracrRNA template, and filled up to 50 μL total volume using molecular biology grade water ^10^. The thermal cycle included an initial denaturation step at 98 ℃ for 10 seconds, 5 PCR cycles for crRNA template or 12 PCR cycles for tracrRNA template with 98 ℃ denaturation for 5 seconds and annealing at 55 ℃ for 15 seconds, and a final extension step at 72 ℃ for 1 minute. The PCR product was directly added to TranscriptAid T7 High Yield Transcription Kit (Thermo Fisher) reaction as the DNA template for RNA transcription following the manufacturer’s instruction, followed by incubating at 37 ℃ for 4-6 hours until significant amount of white mist formed. After transcription, crRNA and tracrRNA were purified by RNA Clean & Concentrator-5 with DNase I Set (Zymo Research Corporation) following the total RNA clean-up protocol with DNase I treatment before RNA clean-up. The DNase I treatment was increased to 30 minutes to ensure thorough removal of residual DNA templates. The 100% ethanol was adjusted to 1.5 volume to increase short-length RNA yield. The RNA product was eluted by 15 μL of DNase/RNase-free water. The purified crRNA and tracrRNA were quantified by Qubit RNA Broad Range Assay Kit (Thermo Fisher) using 100-fold diluted RNA products. The crRNA and tracrRNA were aliquoted and stored separately at −80 ℃ before use ^9^.

#### S4. CRISPR-NGS library preparation

Instead of using non-specific enzymes for DNA fragmentation in most NGS library preparation workflows, in the CRISPR-NGS library preparation workflow, we used Cas9 nuclease to specifically cleave targeted sites for DNA fragmentation. Right before CRISPR-NGS library preparation, crRNA and tracrRNA were mixed equimolarly with the final concentration adjusted by Duplex Buffer (Integrated DNA Technology) to 2,500 ng/μL. The mixture was heated up to 94 ℃ for 2 minutes and slowly cooled down to room temperature for gRNA assembly ^11^. Then the duplexed gRNA was mixed with pre-diluted 25 μM Alt-R™ S.p. HiFi Cas9 Nuclease V3 (Integrated DNA Technologies) in a 10:3 volume ratio and incubated at room temperature for 2.5 hours to form Cas9-gRNA complex.

During Cas9-gRNA complex formation, rAPid Alkaline Phosphatase (Roche) was used to remove the 5’ phosphate groups of the DNA samples. A 10 μL phosphatase reaction included 100-300 ng of DNA samples, 1 μL of rAPid Alkaline Phosphatase Buffer (10x concentrated), and 0.5 μL of rAPid Alkaline Phosphatase (1 U/μL). The mixture was incubated at 37 ℃ for 10 minutes for 5’ phosphate group removal followed by 75 ℃ for 2 minutes for phosphatase inactivation.

The Cas9 cleavage reaction included 3 μL of NEBuffer™ r3.1 (New England Biolabs), 10.4 μL of the Cas9-gRNA complex, 1 μL of 0.01 ng/μL 5’ phosphorylated synthesized external 250-bp DNA spike with no cleavage site in its sequence (5’-ACCCATACAAGGAACCCGGCCAGCACTACGCTCACTACGGCCGGTGGTACGGTGGGCACTCCGGTGAAATGCACGTGCTTGGCATGCCGTCAGGCCGTGAAGTCAAGCGCACCCCGGTGTTCAACATGGACAGCAACAAGATGACCATCCACATCGCCTCGCCGGCGCCGGCATACAGTCTGGGGGGAATTCAAGATGGAGAAGGGCGACGAGGTAATGGCGATCCTGACCTCGACAAGTGGAAGACCTG-3’), 6-8 μL of phosphatase treated DNA sample, and the volume was filled up to 30 μL by molecular biology grade water. The mixture was pipette mixed and incubated at 37 ℃ for 16 hours for thorough cleavage.

After the 16-hr incubation, 2.5 μL of RNase A (10 mg/mL) was directly added to the mixture and incubated at room temperature for 15 minutes. Then, 2.5 μL of Thermolabile Proteinase K (New England Biolabs) was added to each sample, and samples were incubated at 37 ℃ for 15 minutes for protein digestion, and at 55 ℃ for 10 minutes for Thermolabile Proteinase K inactivation. The DNA fragments were dA-tailed by Taq polymerase. A master mix was premade by mixing 2 μL of dATP (100 mM), 5 μL of Taq DNA polymerase (5,000 U/mL), 80 μL of Standard Taq Reaction Buffer (New England Biolabs), and filled the volume up to 100 μL by molecular biology grade water. Five microliters of the dA-tailing master mix were added to each reaction and incubated at 72 ℃ for 20 minutes. After dA-tailing, the cleaved DNA samples were ready for adapter ligation.

We used xGen™ UDI-UMI Adapters (Integrated DNA Technologies) in the adapter ligation step to reduce the bias generated by the PCR step afterward. The adapter ligation reaction included 35 μL of the dA-tailed DNA sample, 2 μL of 100-fold diluted adapter, 30 μL of NEBNext® Ultra™ II Ligation Master Mix (New England Biolabs), and 1 μL of NEBNext® Ligation Enhancer. The reaction was incubated at room temperature for 15 minutes. After adapter ligation, 57 μL of AMPure XP Reagent (Beckman Coulter) was added to the reaction for sample clean up. The adapter-ligated DNA samples were eluted by 15 μL 0.1x TE buffer for PCR amplification.

The 50-μL PCR reaction included 15 μL of adapter-ligated DNA sample, 5 μL of xGen™ Library Amplification Primer Mix (Integrated DNA Technologies), 25 μL of NEBNext® Ultra™ II Q5® Master Mix (New England Biolabs), and 5 μL of molecular biology grade water. The thermal cycle of PCR included an initial denaturation step at 98 ℃ for 30 seconds, 22 cycles of PCR step (98 ℃ for 10 seconds and 65 ℃ for 75 seconds), and a final extension step at 65 ℃ for 5 minutes. The PCR product was cleaned by adding 45 μL of AMPure XP Reagent (Beckman Coulter) into the 50 μL reaction. The cleaned PCR product was eluted by 30 μL 0.1x TE buffer as the CRISPR-NGS library. The libraries were stored at −20 ℃ before sequencing.

### S5. Regular NGS library preparation, quality control, and Illumina sequencing

The regular NGS libraries were prepared using NEBNext® Ultra™ II FS DNA Library Prep Kit for Illumina (New England Biolabs). The same amount of DNA samples as the input of the CRISPR-NGS libraries were used to make the regular NGS libraries. One microliter of 0.01 ng/μL 5’ phosphorylated synthesized external 250-bp DNA spike as mentioned in the above section was added ^12^. The library preparation steps followed manufacturer’s instructions with minor adjustments. In the fragmentation step, the samples were incubated at 37 ℃ for 30 minutes. xGen™ UDI-UMI Adapters (Integrated DNA Technologies) was used for adapter ligation, so that the USER® Enzyme incubation step was skipped. The adapter dilution ratios were adjusted by the DNA input for each library (**Table S4**). Five cycles of PCR were used in the PCR enrichment step. The final library was eluted in 30 μL of 0.1x TE buffer and stored at −20 ℃ before sequencing.

The concentrations of the libraries were determined by Qubit™ 1X dsDNA High Sensitivity (HS) Assay Kit (Invitrogen) right after library preparation. After concentration check, the libraries were sent to Roy J. Carver Biotechnology Center for following quality check steps and sequencing. DNA fragment lengths were determined by AATI Fragment Analyzer. An additional beads purification was conducted before sequencing to remove excess adapter dimers according to the fragment analysis results. A small portion of the libraries were pooled and pre-sequenced on Illumina MiSeq V3 to determine the optimal pooling ratio. Then, all libraries were pooled according to the MiSeq read depths and sequenced on Illumina NovaSeq X Plus 10B paired-reads.

#### S6. Sequencing read mapping

After paired-end sequencing, the UMI indexes were extracted into a separated fastq file. For each library, three fastq files were generated to store read 1, read 2, and the UMI indexes. All sequencing raw data, including the separated UMI index files, can be found in BioProject PRJNA1148079 (to be released upon publication) ^13^. All fastq files were sorted by the command “cat file.fastq | paste | sort -k1,1 -t “ ” | tr “\t” “\n” > file_sorted.fastq” before processing. UMI-Tools was used to add UMI indexes from the separated UMI index file to corresponding identifiers in read 1 and read 2 files using the command “umi_tools extract --extract-method=string --bc-pattern=NNNNNNNNN -I [UMI_index_fastq_file] -S [UMI_index_fastq_file_out_nothing_left] --read2-in=[read_fastq_file_to_add_UMI] read2-out=[output_read_fastq_file_with_UMI_added]” ^14^. After adding UMI indexes, the sequencing reads in read 1 and read 2 files were screened by PriceSeqFilter using the criteria that 85% of nucleotides in a read must be high quality, minimum allowed probability of a nucleotide being correct is 0.98, and 90% of nucleotides in a read that must be called ^15^. The output fastq files from PriceSeqFilter were used for read mapping in Bowtie2 with very sensitive end-to-end mapping and reporting all alignments ^16^. A fasta file including the sequences of all ARGs downloaded from CARD for gRNA design (see 2.3) and the sequence of the 250-bp external DNA spike was used as the reference. The output sam files were converted to bam files by SAMtools using -bSF4 command to remove unaligned reads^17,18^. The bam files were sorted and indexed by SAMtools before being deduplicated by UMI-Tools according to the UMI indexes added to the read identifiers earlier ^14^. The coverage and average read depths of the mapping for all ARGs and the external DNA spike were exported by the coverage function in SAMtools using the deduplicated bam file as the input with “--ff 0” setting to remove the restriction on secondary alignment ^17,18^. Any ARGs with coverages higher than 75% were considered present in the sample, according to previous studies ^19–21^. The identity threshold was approximately 90%, according to the end-to-end alignment algorithm of Bowtie2 ^16^.

After the above steps, a validation step was conducted to check the reliability of read mapping. In all ARG families that were detected by sewage CRISPR-NGS libraries, one ARG from each family was randomly selected and viewed for these CRISPR libraries in Integrative Genomics Viewer (IGV) ^22^. Since DNA in CRISPR-NGS libraries were fragmented by cleaving at designed sites and DNA in regular NGS libraries were fragmented randomly, sharp boundaries of reads were expected at cleavage sites for CRISPR-NGS libraries while the read mapping histograms were smooth for regular NGS libraries (**Figure S1**). Read mapping was checked by comparing the cleavage sites shown in IGV to the bed file generated after gRNA design indicating all possible cleavage sites for all ARGs.

#### S7. Validation for 5’ phosphate group removal

A synthesized 731-bp DNA fragment with 5’ phosphorylation was used for the validation of 5’ phosphate group removal by alkaline phosphatase. The synthesized DNA was dissolved to a concentration of 10 ng/μL by molecular grade water to avoid inhibition to the enzymes by EDTA. The 5’ phosphate group removal reaction (10 μL) included 8.5 μL of synthesized DNA, 1 μL of rAPid Alkaline Phosphatase Buffer (10x concentrated), and 0.5 μL of rAPid Alkaline Phosphatase (1 U/μL). The mixture was incubated at 37 ℃ for 10 minutes for 5’ phosphate group removal followed by 75 ℃ for 2 minutes for phosphatase inactivation. A negative control sample was made by replacing the alkaline phosphatase by the same volume of molecular biology grade water. Then, both samples were used as the DNA input of T4 ligation. The ligation reaction included 8.5 μL of either positive or negative DNA sample, 1 μL of T4 DNA Ligase Buffer (10X), and 0.5 μL of T4 DNA Ligase (New England Biolabs). The mixture was incubated at room temperature for two hours for ligation. Then, the ligated DNA samples were loaded to agarose gel for ligation results visualization. Phosphatase-treated DNA did not have 5’ phosphate groups, so that the DNA fragments would not ligate to form new fragments. This sample would show a single band in gel. In contrast, the DNA sample that was not treated by alkaline phosphatase should still have 5’ phosphate group for ligation. This sample would show multiple bands in gel.

#### S8. ARG clustering for quantification

Among the 4,641 ARGs used for gRNA design, some have highly similar sequences with only a few mutations. One read may be mapped to multiple ARGs that share highly similar sequences. To avoid the bias in ARG quantification due to the overlaps in read mapping, we divided the 4,641 ARGs into 1,208 clusters based on their average nucleotide identities (ANI). The pairwise ANIs for all ARGs were calculated by FastANI using fragment length of 100 bp and minimum fraction of a sequence that must be shared to be 100% ^23^. The FastANI output was screened by a customized python code to keep only the ANIs above 90%. A network was built for the screened ANIs in Cytoscape ^24^. The ARGs were plotted as nodes. Two nodes were connected if the ANI between them was greater than 90%. The connected nodes were defined to be in the same cluster. If a node did not connect to any other nodes, it was in an individual cluster. There were 228 clusters containing more than one ARGs, and 980 clusters containing only one single ARG. The read depth of a cluster was calculated by the average read depths of all ARGs in this cluster.

#### S9. Reproducibility test for CRISPR-NGS libraries

To test the reproducibility of CRISPR-NGS in ARG detection and quantification, the libraries for Calumet (8/16/2023) and Calumet (10/24/2023) were made in duplicates. After sequencing and read mapping, scatter plots were made in log-10 scale using the read depths of the ARGs detected in both duplicated libraries as the x and y coordinates. Then, the scatter plots were fitted linearly to get the correlation of the same ARGs detected in both libraries. OriginPro 2024 was used to make the scatter plots and do the fitting.

#### S10. CRISPR-NGS read subsampling

To compare the ARG detection efficiency between CRISPR-NGS and regular NGS with higher read depth, we subsampled the CRISPR-NGS fastq files by 1%, 2%, 5%, 10%, 20%, 50%, and 70% using seqtk (https://github.com/lh3/seqtk/tree/master) with the same seed -s100 after reads sorting, UMI index labeling, and reads quality filtering as described in section 2.5 in the main manuscript. After subsampling, all subsampled fastq files were mapped to ARGs following the same protocol as described in section 2.5. After getting the ARG mapping results of each individual subsampled CRISPR-NGS library, we calculated the ratio between the number of ARGs detected by subsampled CRISPR-NGS libraries and the corresponding regular NGS libraries. Then, these ARG detection ratios were plotted in OriginPro 2024 against the subsampling ratio.

#### S11. Sanger sequencing for ARG presence validation

Sanger sequencing was conducted to validate the presence of the ARGs detected in the DNA samples. The ARGs targeted in this validation step were randomly selected from the ARG mapping results. The sequences of the selected ARGs were downloaded from CARD ^25^. Then, these sequences were used as the input for Primer BLAST on NCBI to design primers for long-range PCR ^26^. The minimum PCR product size was modified to be less than 100 bp smaller than the size of the input ARG to ensure covering the largest possible segment of the gene. The database used in Primer BLAST was nr, and the organism was set as bacteria (taxid: 2). The long-range PCR primers are listed in **Table S2**.

For long-range PCR, we added 12.5 μL of Phusion® High-Fidelity PCR Master Mix (New England Biolabs) or NEBNext® Ultra™ II Q5® Master Mix (New England BIolabs), 1.25 μL of 10 μM forward and reverse primers respectively, 1 μL of the DNA sample, and filled the volume up to 25 μL by nuclease-free water for each reaction. The PCR thermal cycle includes an initial denaturation step of 98 ℃ for 30 seconds, 34 cycles of the PCR step with 10 seconds of denaturation at 98 ℃, 30 seconds of annealing at 55 ℃, and 90 seconds of elongation at 72 ℃, and a final extension step at 72 ℃ for 5 minutes. The amplicon lengths and DNA concentrations were validated by DNA electrophoresis and Qubit™ 1X dsDNA High Sensitivity (HS) Assay Kit (Thermo Fisher) before Sanger sequencing. Sanger sequencing was conducted by UIUC Core Sequencing Facility. Each DNA sequence was aligned to the corresponding reference gene sequence in CARD by NCBI nucleotide BLAST to test their similarity ^8^.

#### S12. qPCR for ARG quantification validation

Nine ARGs with various read depths detected by CRISPR-NGS or regular NGS methods were selected for qPCR validation. The primers for these ARGs and their corresponding amplification efficiencies are listed in **Table S5**. A 15-μL qPCR reaction included 7.5 μL of PowerUp SYBR Master Mix (Applied Biosystems), 600 nM of forward and reverse primers, 2 μL of 10-fold diluted sewage DNA extract, and 4.3 μL of molecular biology grade water. The samples were loaded in triplicates in a qPCR run. Reactions targeting the 16S rRNA gene for the same samples were also loaded in triplicates on every qPCR plate for relative abundance quantification. qPCR was performed on QuantStudio 7 Pro (Thermo Fisher) using fast cycle followed by a melt curve stage for the quality control of non-specific amplification. The qPCR thermal cycle included an initial hold stage at 95 ℃ for 20 seconds and 40 PCR cycles with 95 ℃ for 1 second and 60 ℃ for 20 seconds. After the qPCR run, the relative abundances of the ARGs were determined by ΔCq method using the following equation ^27^:

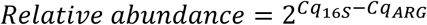

In the equation, Cq_16S_ means the average Cq value of the 16S rRNA gene in the sample, and Cq_ARG_ means the average Cq value of the ARG in the same sample. After obtaining the qPCR results for the nine ARGs in all sewage samples, the read depths in both CRISPR-NGS and regular NGS against the relative abundances in qPCR were plotted and fitted as linear regression curves in log10 scale in Origin Pro 2024 to evaluate the ARG quantification potential for the CRISPR-NGS method.

### Supplementary Tables

**Table S1.** Mock community composition.

| Isolate | Species/Genus |
| --- | --- |
| Environmental E. coli 1 | *E. coli* |
| Environmental E. coli 2 | *E. coli* |
| Environmental E. coli 3 | *E. coli* |
| Clinical E. coli 1 | *E. coli* |
| Clinical E. coli 2 | *E. coli* |
| Clinical E. coli 3 | *E. coli* |
| Clinical Shigella 1 | *Shigella* spp. |
| Clinical Salmonella 1 | *Salmonella* spp. |
| Clinical Salmonella 2 | *Salmonella* spp. |
| PA14 | *Pseudomonas aeruginosa* |

**Table S2.**
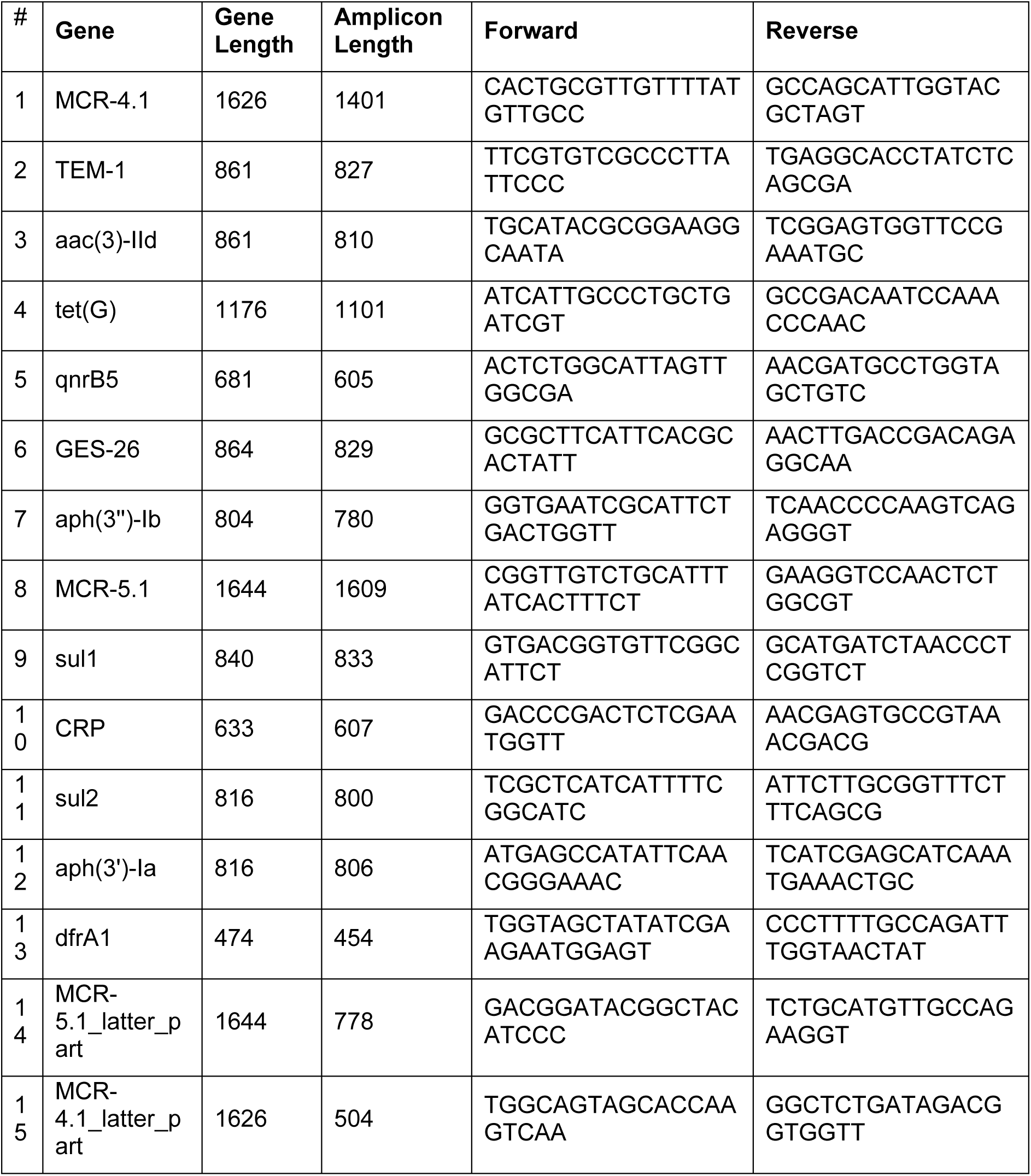
The PCR primers designed for Sanger sequencing validation.

**Table S3.** Summary of positive and negative results in ARG detection using CRISPR-NGS method for the mock community sample.

| ARG cluster | True | False |
| --- | --- | --- |
| **Positive** | 123<br>(97 from whole-genome sequencing, 25 from BLAST, 1 from Sanger sequencing) | 1 (*aph(3')-Ia*) |
| **Negative** | NA | 2 (identity < 90% in whole-genome sequencing annotation) |

**Table S4.** Information for library preparation and sequencing.

| Method | Library | Adapter dilution ratio | DNA input (ng) | NH8B spike amount (ng) | Total number of reads |
| --- | --- | --- | --- | --- | --- |
| CRISPR-NGS | Mock community | 1:100 | 123.8 | 8.66E-03 | 68,633,041 |
|  | Calumet 8/16/2023 | 1:100 | 108.0 | 5.40E-02 | 58,184,677 |
|  | Calumet 8/16/2023 | 1:100 | 115.2 | 8.06E-03 | 47,361,080 |
|  | Calumet 10/24/2023 | 1:100 | 111.6 | 4.40E-03 | 49,296,820 |
|  | Calumet 10/24/2023 | 1:100 | 111.6 | 4.40E-03 | 43,305,745 |
|  | Calumet 12/19/2023 | 1:100 | 110.4 | 6.86E-03 | 52,492,274 |
|  | Lemont 8/16/2023 | 1:100 | 121.2 | 6.86E-03 | 51,487,769 |
|  | Lemont 12/19/2023 | 1:100 | 136.8 | 6.86E-03 | 50,975,555 |
|  | Campus Town 11/30/2021 | 1:100 | 65.8 | 8.06E-03 | 54,097,785 |
| Regular NGS | Mock community | 1:1 | 122.9 | 7.60E-03 | 58,886,378 |
|  | Calumet 8/16/2023 | 1:1 | 108.0 | 1.11E-01 | 49,880,963 |
|  | Calumet 10/24/2023 | 1:1 | 178.5 | 8.40E-03 | 57,076,379 |
|  | Calumet 10/24/2023 | 1:1 | 178.5 | 8.40E-03 | 44,601,769 |
|  | Calumet 12/19/2023 | 1:1 | 166.4 | 7.60E-03 | 57,647,030 |
|  | Lemont 8/16/2023 | 1:1 | 161.2 | 7.60E-03 | 60,683,061 |
|  | Lemont<br>12/19/2023 | 1:1 | 200.6 | 7.60E-03 | 51,761,671 |
|  | Campus Town<br>11/30/2021 | 1:10 | 100.1 | 7.60E-03 | 46,173,438 |

**Table S5.**
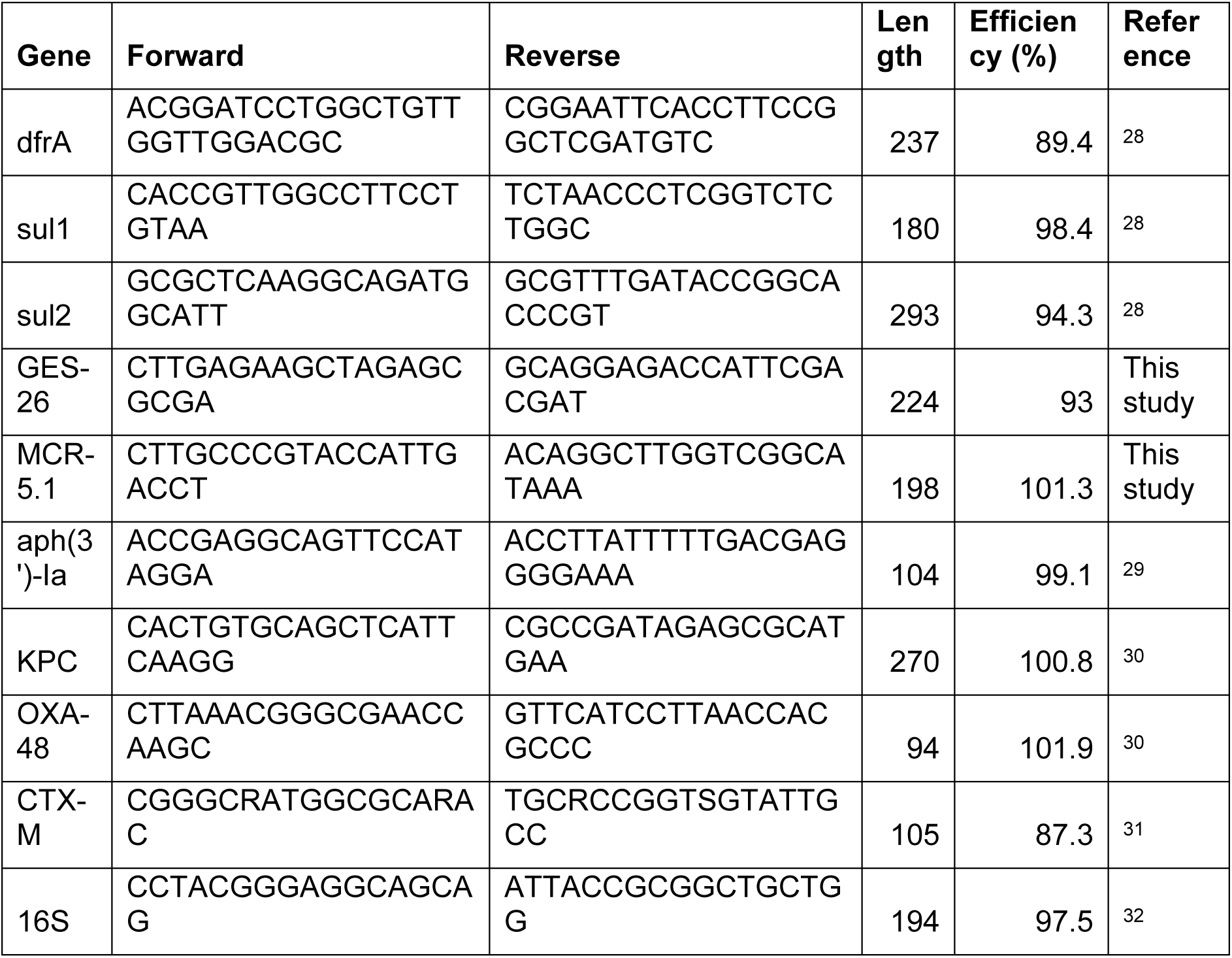
Information for primers used for qPCR.

**Table S6.**
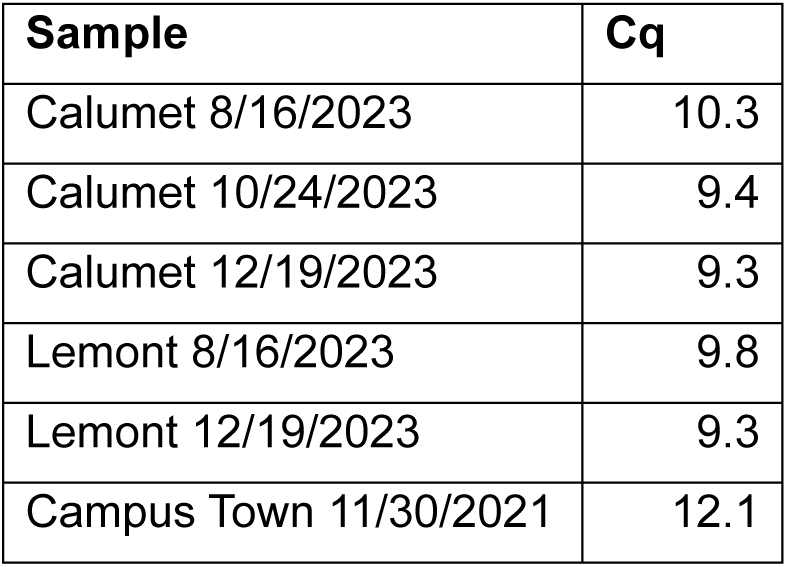
The Cq values of the 16S rRNA gene in the six sewage samples.

**Table S7.** All sewage samples used in this study.

| Sample | Time | Location | Description | Volume (mL) |
| --- | --- | --- | --- | --- |
| Calumet WWTP 1 | 8/16/2023 | Chicago | Wastewater influent | 50 |
| Calumet WWTP 2 | 10/24/2023 | Chicago | Wastewater influent | 50 |
| Calumet WWTP 3 | 12/19/2023 | Chicago | Wastewater influent | 50 |
| Lemont WWTP 1 | 8/16/2023 | Chicago | Wastewater influent | 75 |
| Lemont WWTP 2 | 12/19/2023 | Chicago | Wastewater influent | 50 |
| Campus Town manhole | 11/30/2021 | Central Illinois | Raw sewage from a manhole | 20 |

### Supplementary Figures

**Figure S1.**
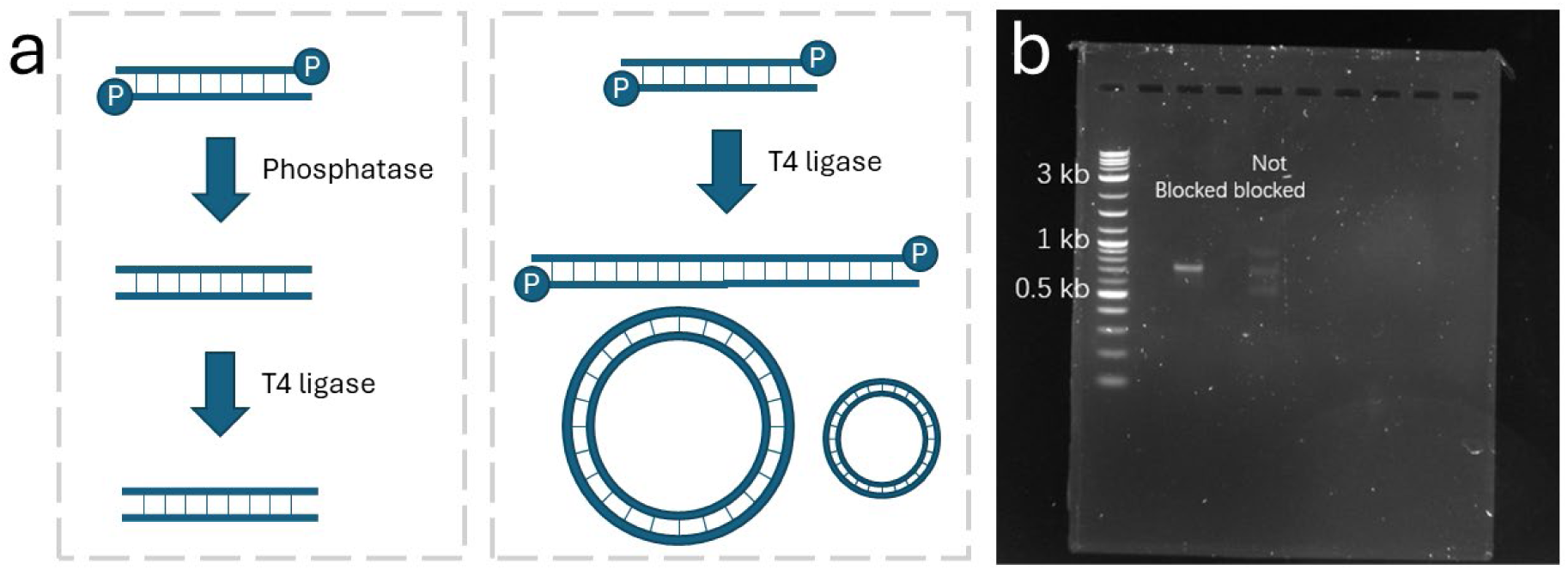
Validation of alkaline phosphatase blocking to remove 5’ phosphate. a: The concept figure of 5’ phosphate removal validation using T4 ligase. b: The gel image of blocked and unblocked samples after T4 ligase treatment.

**Figure S2.**
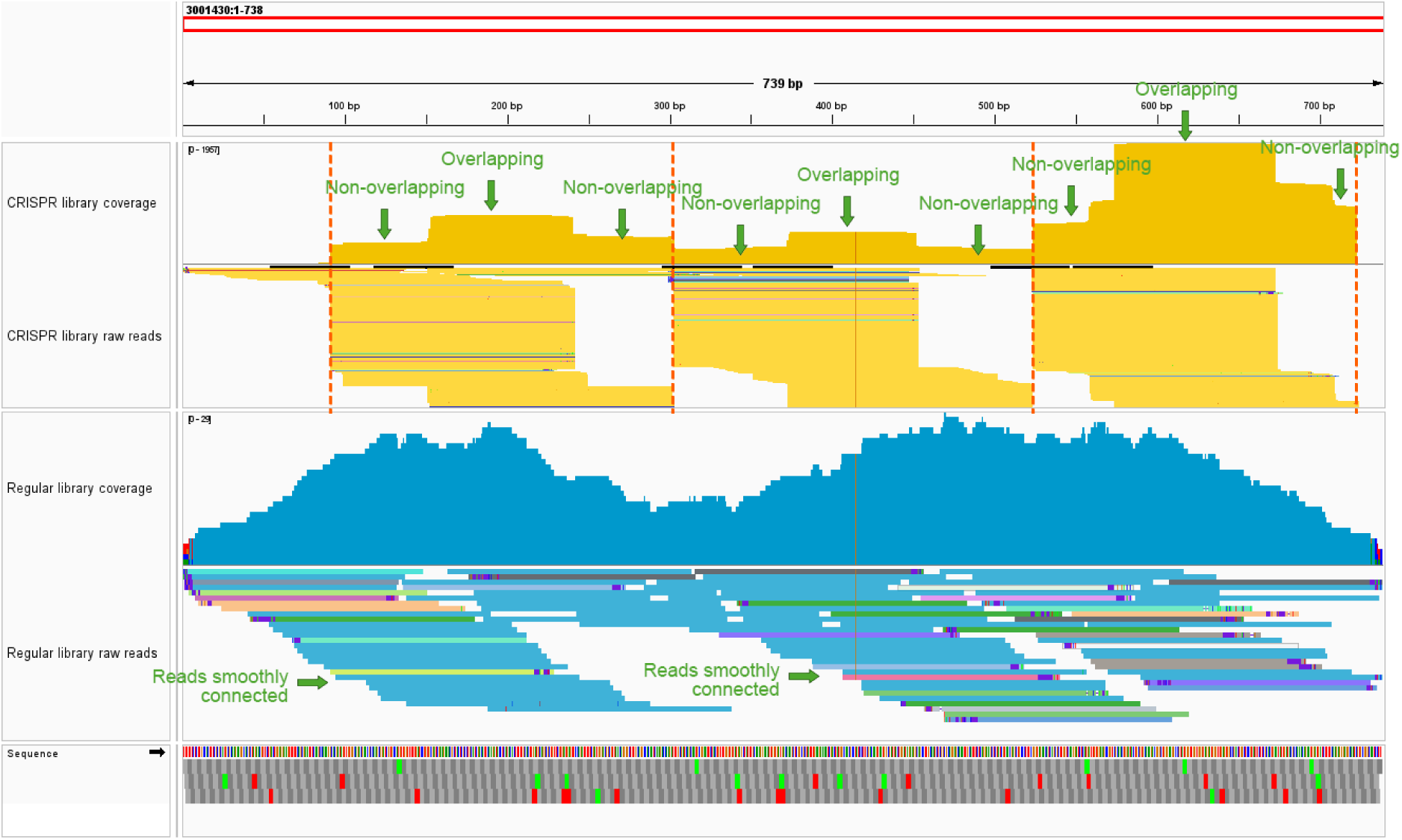
The difference in read mapping between the CRISPR-NGS library (top) and the regular NGS library (bottom) using the example of OXA-36 (ARO: 3001430). The number axis at the top of the figure shows the scale of OXA-36 in base pairs (bp). The DNA fragments were cleaved at the four gRNA binding sites (85-108 bp, 296-319 bp, 517-540 bp, and 706-729 bp, labeled by orange dash lines) for the CRISPR-NGS library.

**Figure S3.**
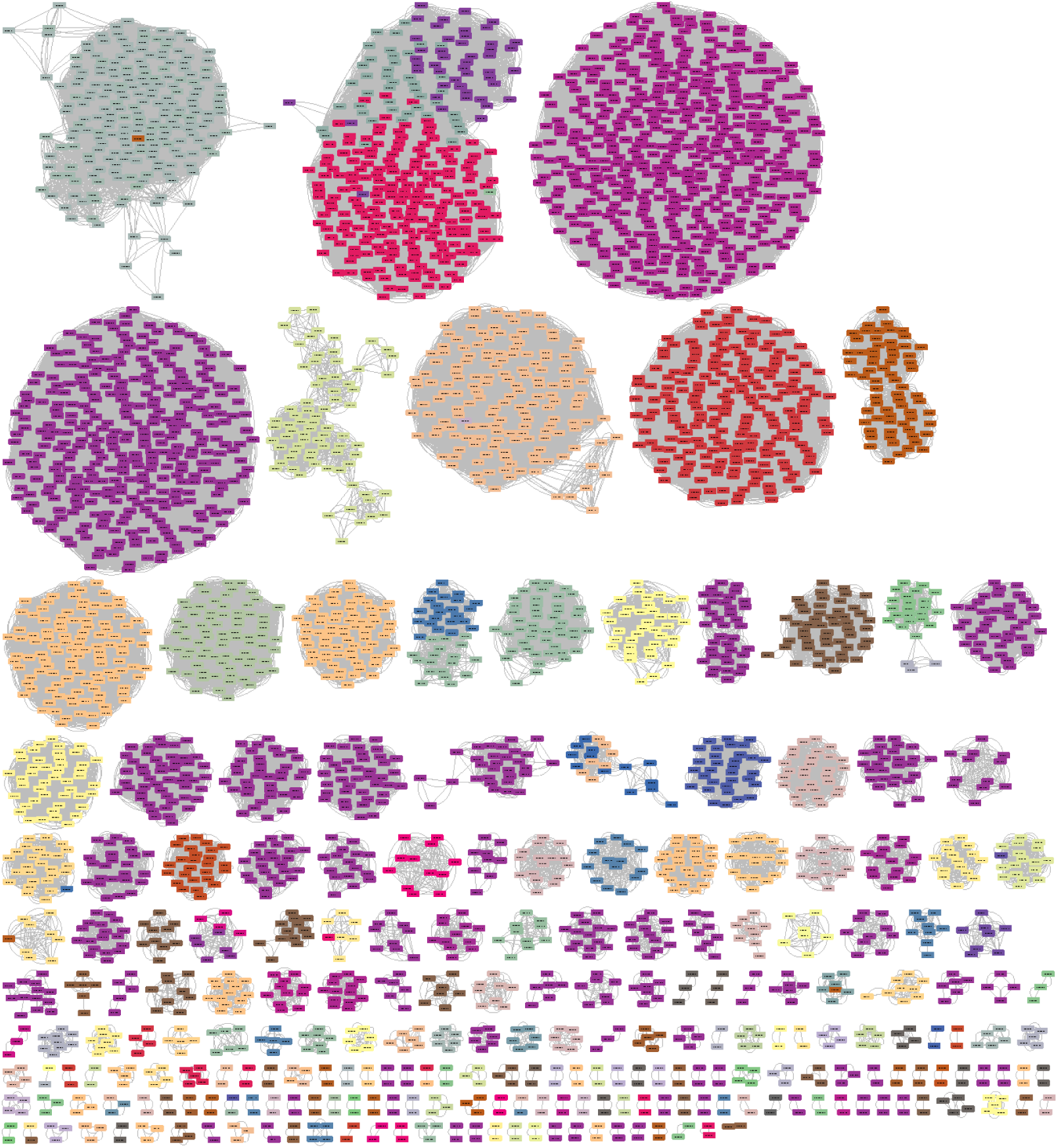
ARG clusters containing more than one ARGs.

**Figure S4.**
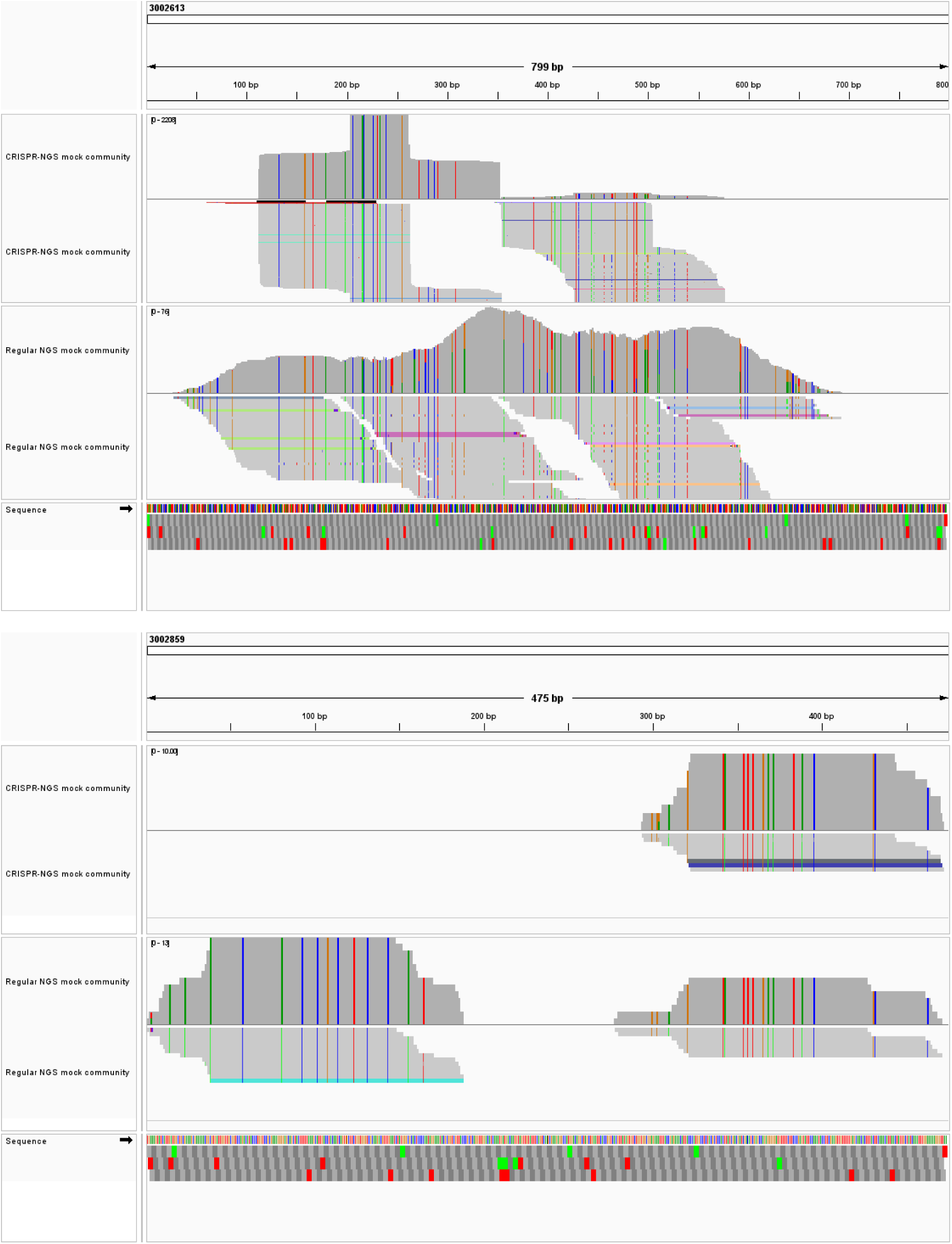
The comparison of read mapping of *aadA13* (ARO: 3002613, 799 bp) (top) and *dfrA14* (ARO: 3002859, 475 bp) (bottom) in the mock community sample for the CRISPR-NGS library and the regular NGS library.

**Figure S5.**
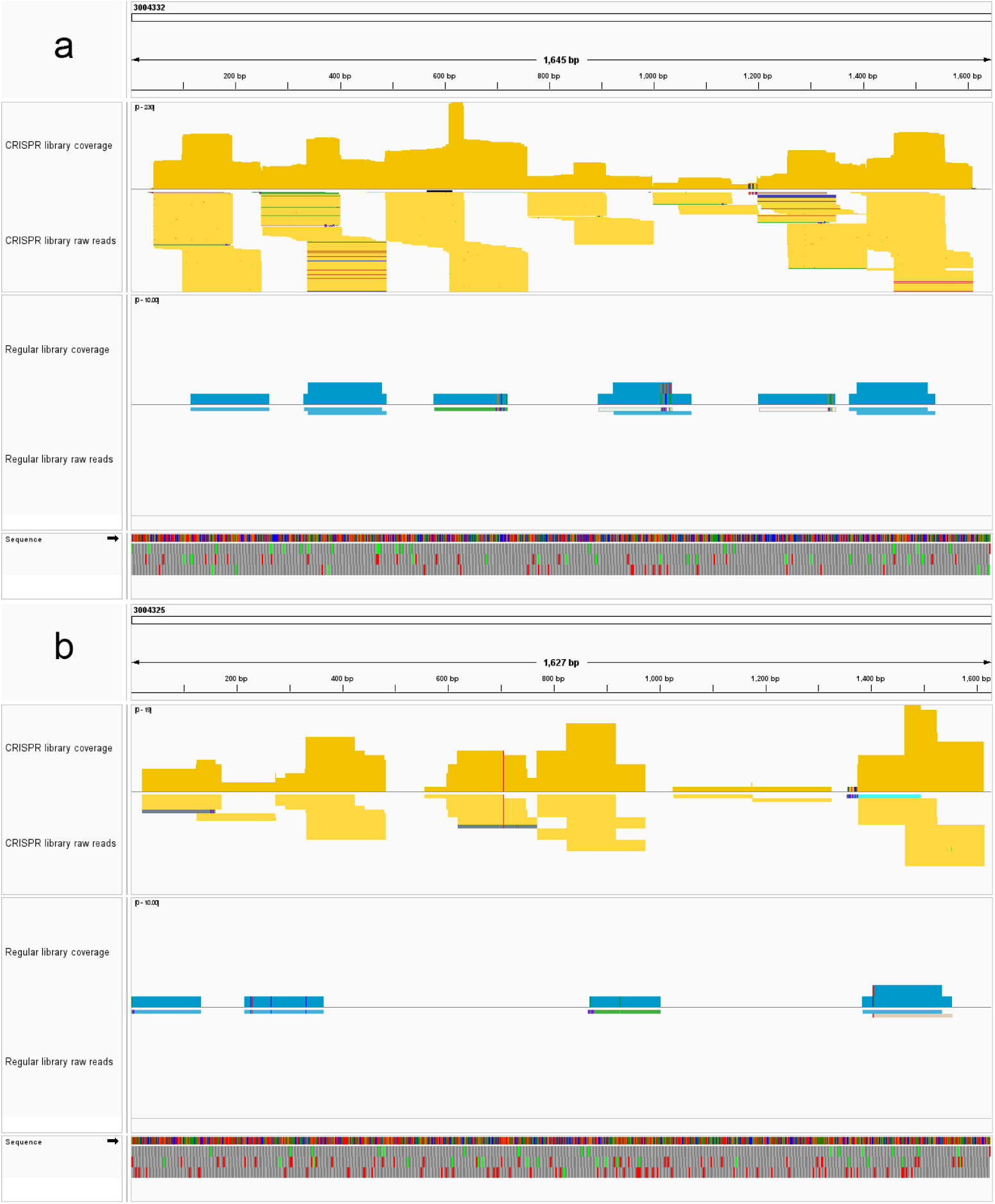
The read mapping of MCR-5.1 (ARO: 3004332) (a) and MCR-4.1 (ARO: 3004325) (b) in the CRISPR-NGS library (yellow) and the regular NGS library (blue) of Lemont (8/16/2023) sample.

**Figure S6.**
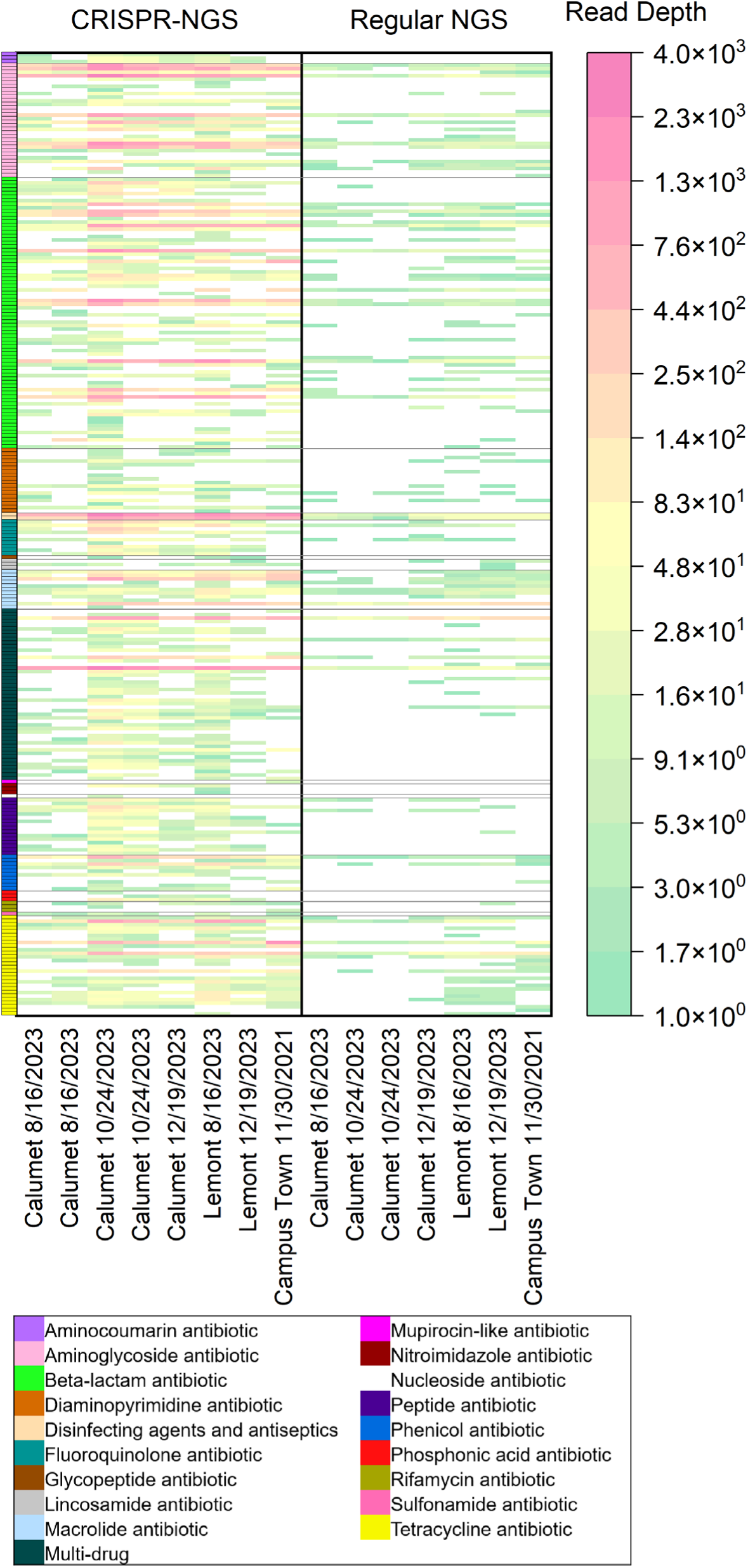
The heatmap for the average read depths of all 270 ARG clusters in all sewage samples measured by CRISPR-NGS method and regular NGS method. The read depths are colored in log-10 scale. The read depths increase from green to pink. The ARG clusters are sorted by different drug classes. The drug classes are labeled by different colors on the left.

### Supplementary Spreadsheets

**Spreadsheet S1.** The clustering of different ARGs targeted in this study.

**Spreadsheet S2.** The ARGs detected in the mock community sample by CRISPR-NGS.

**Spreadsheet S3.** The ARG annotation results for the 10 PacBio HiFi whole-genome sequences used in mock community.

**Spreadsheet S4.** The ARGs detected in the mock community sample by regular NGS.

**Spreadsheet S5.** The ARGs detected in all sewage samples by CRISPR-NGS.

**Spreadsheet S6.** The ARGs detected in all sewage samples by regular NGS.

**Spreadsheet S7.** The gRNA spacer sequences designed in this study.

